# BSCL2/Seipin Deficiency in Heart Causes Energy Deficit and Heart Failure via Inducing Excessive Lipid Catabolism

**DOI:** 10.1101/2021.06.07.447355

**Authors:** Hongyi Zhou, Jie Li, Huabo Su, Ji Li, Todd A. Lydic, Martin E Young, Weiqin Chen

## Abstract

Heart failure (HF) is one of the leading causes of death world-wide and is associated with cardiac metabolic perturbations. Human Type 2 Berardinelli-Seip Congenital Lipodystrophy (BSCL2) disease is caused by mutations in the *BSCL2* gene. Global lipodystrophic *Bscl2^−/−^* mice exhibit hypertrophic cardiomyopathy. Whether BSCL2 plays a direct role in regulating cardiac substrate metabolism and/or contractile function remains unknown. Here we show that mice with cardiac-specific deletion of *Bscl2* (*Bscl2^cKO^*) developed dilated HF. Myocardial BSCL2 deletion led to elevated ATGL expression and FA oxidation (FAO) along with reduced cardiac lipid contents. Cardiac dysfunction in *Bscl2^cKO^* mice was independent of mitochondrial dysfunction and oxidative stress, but associated with decreased metabolic reserve and ATP levels. Importantly, heart failure in *Bscl2^cKO^* mice could be partially reversed by pharmacological inhibition of FAO, or prevented by high fat diet (HFD) feeding. Lipidomic analysis further identified markedly reduced glycerolipids, glycerophospholipids, NEFA and acylcarnitines in *Bscl2^cKO^* hearts, which were partially normalized by FAO inhibition or HFD. Our study reveals a new form of HF with excessive lipid catabolism, and identifies a crucial cardiomyocyte-specific role of BSCL2 in controlling cardiac lipid catabolism, energy state and contractile function. It also provides novel insights into metabolically treating energy-starved HF using FAO inhibitor or HFD.

## Introduction

Heart failure (HF) is one of the leading causes of morbidity and mortality worldwide^1^. In healthy individuals, the heart exhibits striking metabolic flexibility, being capable of utilizing carbohydrate, lipid, amino acids and/or ketone bodies to meet energetic demands, cellular constituent turnover, and metabolic signaling; these processes are critical for maintenance of mechanical work. Oxidation of fatty acids (FAs) predominates, accounting for 60-70% of myocardial oxygen consumption^2, 3^. In patients with hypertrophied and failing hearts, derangements of substrate utilization include an increased reliance on glycolysis concomitant with an overall reduced oxidative metabolism^4^. The severely failing heart usually demonstrates a lower concentration of ATP, supporting the concept that energy starvation contributes significantly to the pathogenesis of HF^5^. Indeed, there is a striking correlation between cardiac energetic status and survival in HF patients^6^. Thus, targeting metabolic processes in the heart may represent a promising way to develop new therapeutic approaches for HF.

Cardiac lipid metabolism is precisely controlled to maintain a balance between FA uptake, triglyceride (TG) synthesis, TG hydrolysis, and FA oxidation (FAO)^7^. Imbalances in these processes are commonly seen in obese and diabetic patients (as well as animal models), which are associated with cardiac steatosis and contractile dysfunction^7^. Recent preclinical and clinical evidence argues for an important role of adipose triglyceride lipase (ATGL)-mediated cardiac lipolysis in promoting mitochondrial FAO and ATP production thus contractile function^8^. Constitutive *Atgl^−/−^* mice develop severe cardiac steatosis and HF, associated with a high mortality^9^. Conversely, cardiomyocyte-specific overexpression of ATGL maintains normal cardiac function in lean mice, and reduces cardiac TG content and improves cardiac function during diabetes and obesity^10, 11^. The precise mechanisms linking cardiac lipid accumulation with contractile dysfunction remain obscure.

Berardinelli-Seip Congenital Lipodystrophy 2 (BSCL2, a.k.a. SEIPIN) is a highly conserved endoplasmic reticulum (ER) protein expressed in most tissues, with the highest levels in testis, neuronal and adipose tissue^12^. Global *Bscl2*-deficient (*Bscl2^−/−^*) mice recapitulate human BSCL2 disease, exhibiting congenital lipodystrophy and severe insulin resistance^13–15^. Various molecular functions of BSCL2 have been proposed, ranging from regulating lipid droplet (LD) biogenesis^16, 17^ to mitochondrial metabolism^18, 19^. Human and *Drosophila* BSCL2 assembles as an undecamer or dodecamer respectively and play crucial roles in lipid transfer and/or LD formation^20, 21^. BSCL2 has been shown to interact with 1-acylglycerol-3-phosphate O-acyltransferase 2 (AGPAT2)^22^, LIPIN1^23^, glycerol-3-phosphate acyltransferase 3 (GPAT3)^24^ and Promethin^25, 26^. We and others demonstrate that BSCL2 plays a key role in regulating cyclic AMP/protein kinase A (cAMP/PKA) mediated TG lipolysis essential for adipocyte differentiation and maintenance^13, 15^. Recently, we reported that *Bscl2^−/−^* mice develop hypertrophic cardiomyopathy associated with reduced cardiac steatosis^27^. Despite the relatively low expression of BSCL2 in murine hearts^28^, whether BSCL2 plays a cell-autonomous role in modulating cardiac lipid metabolism and function has not been fully addressed.

The present study highlights BSCL2 as a key player governing cardiac lipid metabolism. Cardiomyocyte-specific deletion of BSCL2 enhanced ATGL expression and FAO, resulting in markedly reduced cardiac lipid reserve, associated with compromised ATP production and contractile dysfunction. Inhibition of FAO or supplying FAs by high fat diet (HFD) feeding partially alleviated cardiac energetic stress and augmented contractility. These findings improve our understanding of how perturbations in lipid utilization/storage contribute towards HF development.

## Results

### Myocardial deletion of BSCL2 induces dilated heart failure

Previously we demonstrated lipodystrophic *Bscl2^−/−^* mice develop cardiac dysfunction ^27^. In order to interrogate the specific role of cardiac BSCL2, we generated a mouse model with a cardiomyocyte-specific deletion of BSCL2 (*Bscl2^cKO^*) via Myh6-Cre. Gene expression analysis confirmed an approximate 75% reduction of *Bscl2* specifically in cardiac muscle but not in liver and skeletal muscle of *Bscl2^cKO^* mice (Supplemental Figure S1A). Current available antibodies were not sensitive enough to detect endogenous BSCL2 protein in murine heart tissue, preventing us from confirming the cardiac deletion of BSCL2 at the protein level.

We next determined the impact of cardiomyocyte-specific deletion of *Bscl2* on cardiac function, in comparison to two distinct control groups [*Cre-; Bscl2^f/f^* (designated as Ctrl) and *Cre+; Bscl2^w/w^*]. No changes were observed in total body weights at either 3 months or 6 months of age between experimental groups (Supplemental Table S1). At 3 months old, there were also no significant differences in ventricle mass in proportion to body weight (Figure 1A) or tibia length (Supplemental Figure S1B). By echocardiography, 3-month-old mice did not display noticeable changes in left ventricular post wall thickness at end systole (LVPWs) (Figure 1B), left ventricular anterior thickness at end systole (LVAWs) (Figure 1C), left ventricular internal diameters at end systole (LVIDs) (Figure 1D), ejection fraction (Figure 1E) and fractional shortening (Figure 1F) between the three experimental groups. When mice aged to 6 months old, ratios of ventricle mass to body weight (Figure 1A) and tibia length (Supplemental Figure S1B) were still comparable in all groups, suggesting no gross cardiac hypertrophy. In agreement with the previous report^29^, 6-month-old *Cre+, Bscl2^w/w^* mice demonstrated no differences in LVPWs, LVAWs, or LVIDs (Figure 1B-D), but exhibited a minor reduction in ejection fraction and fractional shortening (Figure 1E-F) compared with Ctrl mice. In contrast, 6-month-old *Bscl2^cKO^* mice exhibited decreased wall thickness and increased dilation at both systoles and diastoles (Figure 1B-D and Supplemental Table S1), along with marked reductions in contractility compared with Ctrl and *Cre+, Bscl2^w/w^* mice (Figure 1E-G). A 4 chamber analysis confirmed mild dilatation in 6-month-old *Bscl2^cKO^* mice (Figure 1H). Histological analysis of ventricular cross-sectional area showed no evidence of abnormal cardiomyocyte morphology compared with both control groups (Supplemental Figure S1C). In addition, despite similar induction of atrium natriuretic peptide (*Nppa*) in hearts of *Cre+; Bscl2^w/w^* and *Bscl2^cKO^* mice when compared with Ctrl mice, we observed a greater induction of brain natriuretic peptide (*Nppb*) and growth differentiation factor 15 (*Gdf15*), biomarkers for stressed myocardium, in 6-month-old *Bscl2^cKO^* hearts versus Ctrl hearts (Figure 1I). The expression of adult *Myh6* gene expression in *Bscl2^cKO^* hearts was further downregulated relative to Ctrl hearts (Figure 1I). Moreover, the BSCL2 deletion-induced cardiac dysfunction was not accompanied by excessive myocardial fibrosis (assessed by trichrome staining of collagen deposition) in hearts of *Bscl2^cKO^* mice (Supplemental Figure S1D). Together, these data suggest that cardiomyocyte-specific BSCL2 deletion leads to dilated heart failure independent of the long-term expression of transgene Myh6-Cre.

**Figure 1.**
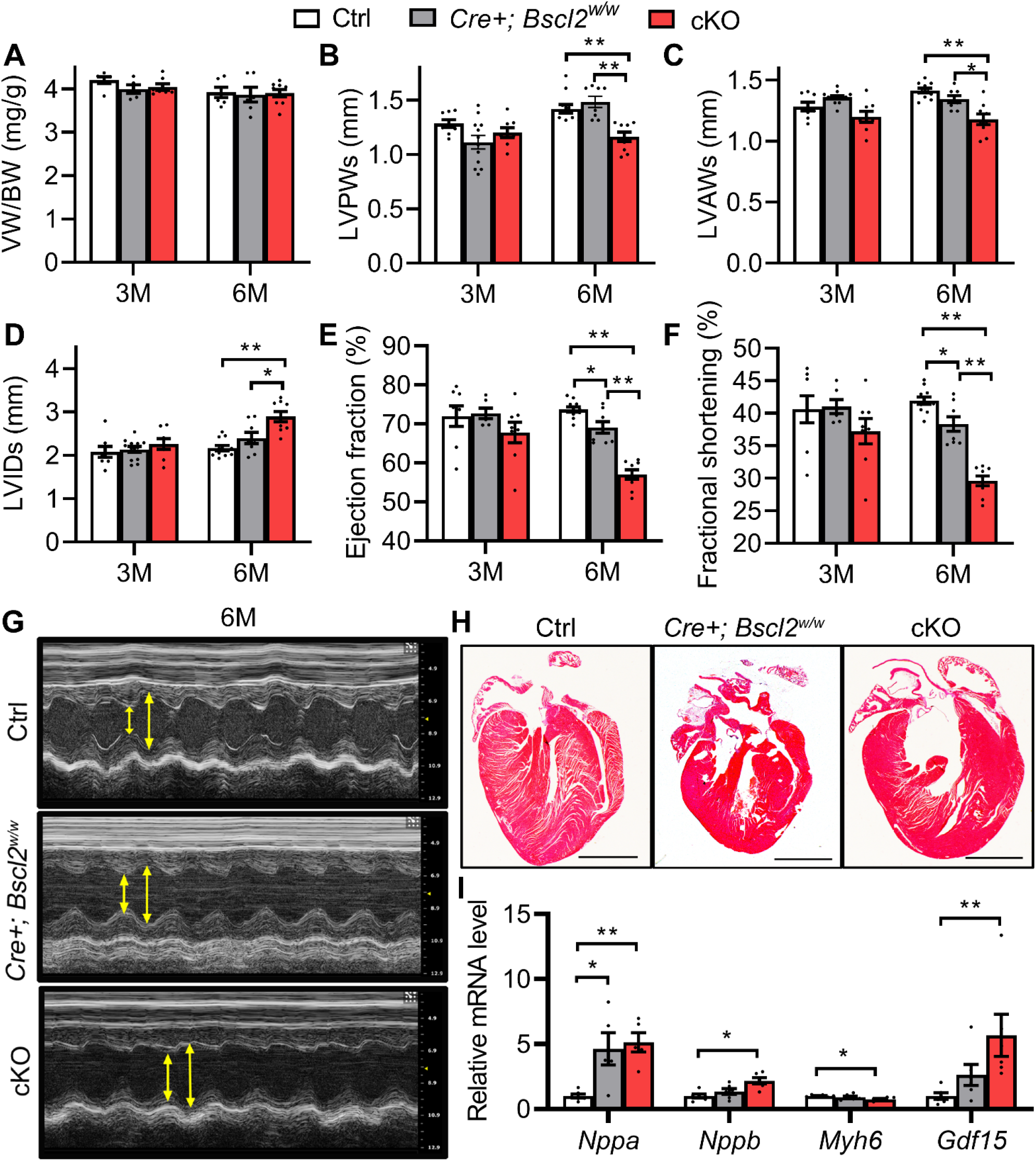
Mice with cardiac-specific deletion of BSCL2 develop dilated cardiomyopathy. **(A)** Ventricle weight (VW) normalized to body weight (BW) in 3-month-old (3M) and 6-month-old (6M) male *Cre-*; *Bscl2^f/f^* (Ctrl), *Cre+*; *Bscl2^w/w^*, and *Cre+*; *Bscl2^f/f^* (cKO) mice. 3M old: Ctrl, *n*=6; *Cre+*; *Bscl2^w/w^*, *n*=5; cKO, *n*=6. 6M old: Ctrl, *n*=6; *Cre+*; *Bscl2^w/w^*, *n*=6; cKO, *n*=9. **(B)** left ventricle post wall thickness at end systole (LVPWs, mm); **(C)** left ventricle anterior wall thickness at end systole (LVAWs, mm); **(D)** left ventricle internal diameter at end systole (LVIDs, mm); **(E)** ejection fraction (%) and **(F)** fractional shortening (%) in 3M and 6M old male mice. 3M old: Ctrl, *n*=8; *Cre+*; *Bscl2^w/w^*, *n*=8; cKO, *n*=12. 6M old: Ctrl, *n*=11; *Cre+*; *Bscl2^w/w^*, *n*=8; cKO, *n*=9. **(G)** Representative echocardiography and **(H)**4 chamber view of 6-month-old mice. Scale bar: 2 mm. **(I)** RT-PCR analysis of cardiac stress genes in ventricles of 6-month-old male mice. *n*=6 per group. *: *P*< 0.05; **: *P*< 0.005. Two-way ANOVA followed by Tukey’s post-hoc tests.

### Myocardial-specific deletion of BSCL2 causes elevated TG turnover and FAO preceding functional decline

Although we have previously demonstrated reduced cardiac TG content, increased ATGL and FAO in lipodystrophic *Bscl2^−/−^* hearts^27^, it remains to be determined whether this was due to a direct loss of BSCL2 from cardiomyocytes. Indeed, we found ventricular TG content in 3-month-old *Bscl2^cKO^* mice was reduced by 57% as compared to Ctrl mice (Figure 2A). *Bscl2^cKO^* hearts displayed increased (≈2.5-fold) ATGL, but not HSL, protein expression compared with that of Ctrl hearts (Figure 2B-C). Such changes were not attributed to alterations in the transcript level of *Pnpla2* (Supplemental Figure S2A), supporting its post-transcriptional regulation. Isolated adult *Bscl2^cKO^* cardiomyocytes displayed similar upregulation of ATGL, but not HSL, compared with Ctrl cells (Figure 2D-E), further supporting a cell-autonomous effect of BSCL2 deletion on ATGL expression. Ctrl and *Bscl2^cKO^* cardiomyocytes responded similarly to the stimulation of isoproterenol in terms of PKA-mediated phospholamban (PL) phosphorylation (Supplemental Figure S2B). Analysis of whole heart lysates also revealed comparable basal PKA-mediated substrate phosphorylation between Ctrl and *Bscl2^cKO^* mice (Supplemental Figure S2C). These data suggest that cardiac-specific deletion of BSCL2 did not affect cAMP/PKA signaling in hearts, different from what we previously observed in BSCL2-deleted adipose tissue^30–32^.

**Figure 2.**
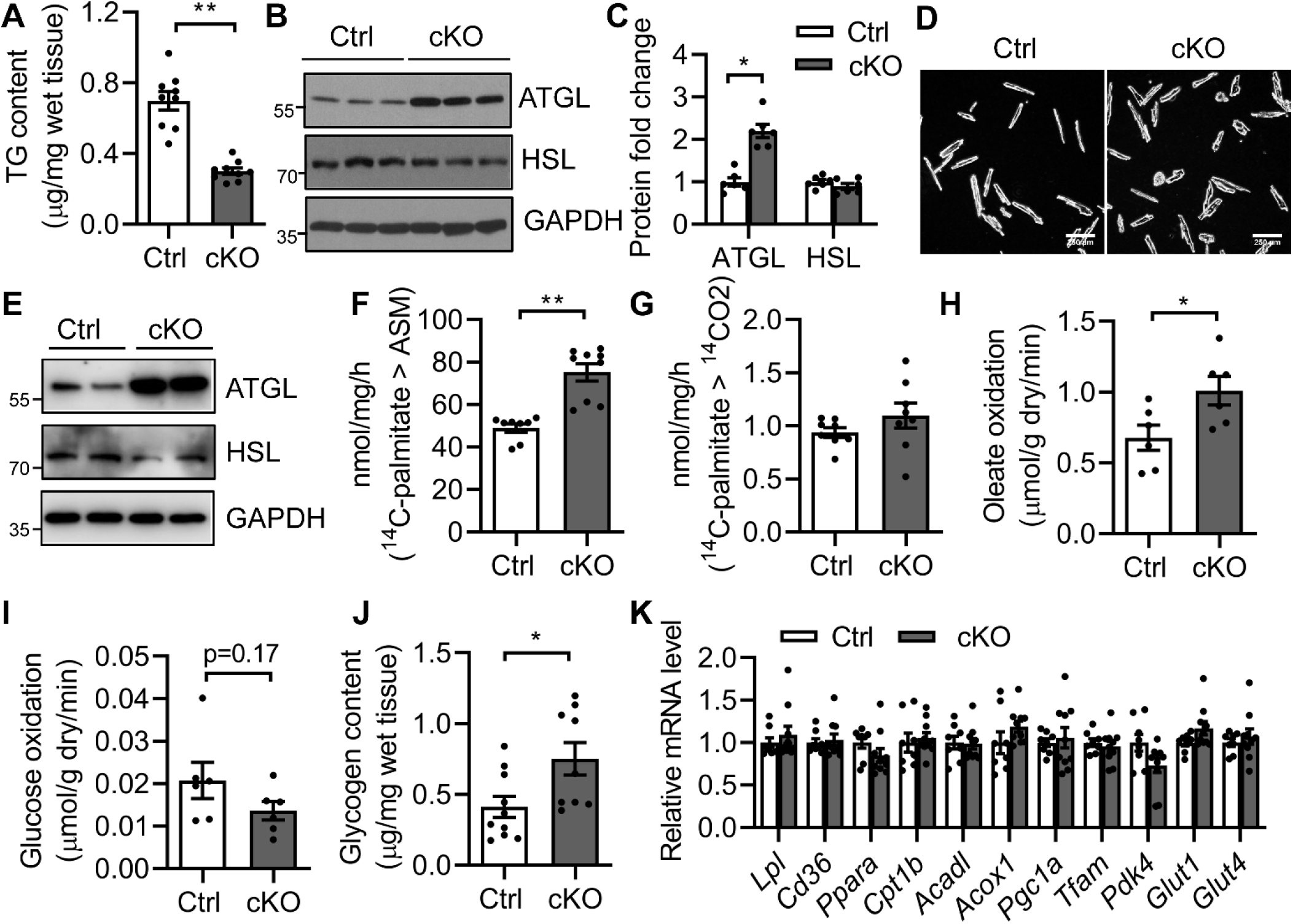
Cardiac-specific deletion of BSCL2 induces cardiac triglyceride turnover and excessive fatty acid oxidation. **(A)** Ventricular TG content as normalized to wet tissue weight. *n*=9 per group. **(B-C)** Representative Western blotting and quantification of lipolytic proteins in heart homogenates. *n*=6 per group. **(D-E)** Isolated adult cardiomyocytes and representative Western blotting in duplicates from two independent experiments. **(F-G)** Acid soluble metabolites (ASM) and CO2 production after incubating heart crude mitochondrial fraction with ^14^C-palmitate. Ctrl, *n*=8; cKO, *n*=9. **(H)** Oleate oxidation rate; **(I)** glucose oxidation rate in *ex vivo* perfused working hearts. *n*=6 per group. **(J)** Glycogen content. Ctrl, *n*=10; cKO, *n*=9. **(K)** RT-PCR analysis of genes involved in fatty acid metabolism, mitochondrial biogenesis and glucose metabolism. Ctrl, *n*=8; cKO, *n*=10. All experiments used 3-month-old male *Cre-*; *Bscl2^f/f^* (Ctrl) and *Bscl2^cKO^* (cKO) mice. *: *P*< 0.05; **: *P*<0.005, unpaired t tests (parametric).

We next investigated FAO in *Bscl2^cKO^* hearts. When incubating heart homogenates with ^14^C-palmitate, 3-month-old *Bscl2^cKO^* mice demonstrated elevated release of ^14^C-labeled acid soluble metabolites (ASM) (Figure 2F) despite a lack of change in ^14^CO_2_ production (Figure 2G). To systemically assess cardiac substrate metabolism preceding cardiac functional decline, hearts from 3-month-old male Ctrl and *Bscl2^cKO^* mice were subject to *ex vivo* perfusions in working mode using radiolabeled substrates. We found no differences in heart rates between two genotypes (Supplemental Figure S2D). The *Bscl2^cKO^* hearts demonstrated a 33% increase in oleate oxidation (Figure 2H) concomitant with a tendency of lower glucose oxidation (Figure 2I). The cardiac oxygen consumption, cardiac efficiency as well as cardiac power (Supplemental Figure S2E-G, respectively) were all comparable between 3-month-old Ctrl and *Bscl2^cKO^* hearts, suggesting maintained cardiac function at this age. Interestingly, the baseline glycogen content in 3-month-old *Bscl2^cKO^* hearts was increased by 83% (Figure 2J), potentially resulting from a tendency toward lower glucose oxidation rate in *Bscl2^cKO^* hearts. In spite of enhanced FAO and glycogen accumulation, *Bscl2^cKO^* hearts demonstrated no changes in the expression of genes involved in FA uptake (*Lpl, Cd36*), mitochondrial and peroxisomal β-oxidation (*Pparα, Cpt1β, Acadl, Acox1*), mitochondrial biogenesis (*Pgc1α, Tfam*), and glucose metabolism (*Pdk4, Glut1, Glut4*) (Figure 2L). Lack of changes in mitochondrial biogenesis was further confirmed by the similar ratios of mitochondrial DNA (mtDNA)-encoded *mt-Rnr2* to nuclear DNA (nDNA)-encoded *Hk2* intron 9 (Supplemental Figure S2H) and protein levels of each of the electron transport chain (ETC) complexes between heart lysates of 3-month-old Ctrl and *Bscl2^cKO^* mice (Supplemental Figure S2I). Together, these data clearly suggest that cardiomyocyte-specific BSCL2 deficiency results in higher rates of cardiac TG turnover and FAO independent of transcriptional changes of mitochondrial and extra-mitochondrial metabolic genes.

### Chronic derangements in myocardial FAO leads to massive lipid remodeling and reduced endogenous substrates in *Bscl2^cKO^* hearts

To identify mechanisms underlying the progressive development of HF, we performed untargeted lipidomic analyses of ventricles from 6-month-old Ctrl and *Bscl2^cKO^* mice. Total normalized lipid ion abundances identified in *Bscl2^cKO^* hearts were reduced by about 45% (Figure 3A). Heatmap analysis revealed massive reductions in the absolute levels of five broadly classified lipid classes defined by the Lipid MAPS consortium; i.e. glycerophospholipids, fatty acyls [mainly nonesterified fatty acids (NEFA)], sphingolipids, sterol lipids and glycerolipids in *Bscl2^cKO^* mice (Figure 3B and Supplemental Excel 1). When comparing the % distributions of these five lipid classes, the proportions of glycerophospholipids, sphingolipids and sterol lipids were significantly higher in *Bscl2^cKO^* hearts as compared to Ctrl hearts (Figure 3C). The proportions of NEFA were relatively comparable between two genotypes, while the proportion of glycerolipids was markedly reduced by 58% in *Bscl2^cKO^* hearts (Figure 3C). Analysis of the absolute levels of glycerolipids revealed 78%, 50% and 75% reductions in TG, diacylglycerol (DAG) and monoacylglycerol (MAG), respectively in *Bscl2^cKO^* hearts (Figure 3D). The absolute levels of NEFA and total acylcarnitines (ACs) were significantly lower in *Bscl2^cKO^* hearts (Figure 3E). Specifically, the abundances of the major long-chain ACs (AC16:0, AC16:1, AC18:0, AC18:1, AC18:2) were all reduced by approximately 70% (Figure 3F). These data suggest myocardial BSCL2 deletion results in a dramatic remodeling of lipid composition highlighted by reduced levels of energy providing substrates indicative of impairment of cardiac metabolic reserve.

**Figure 3.**
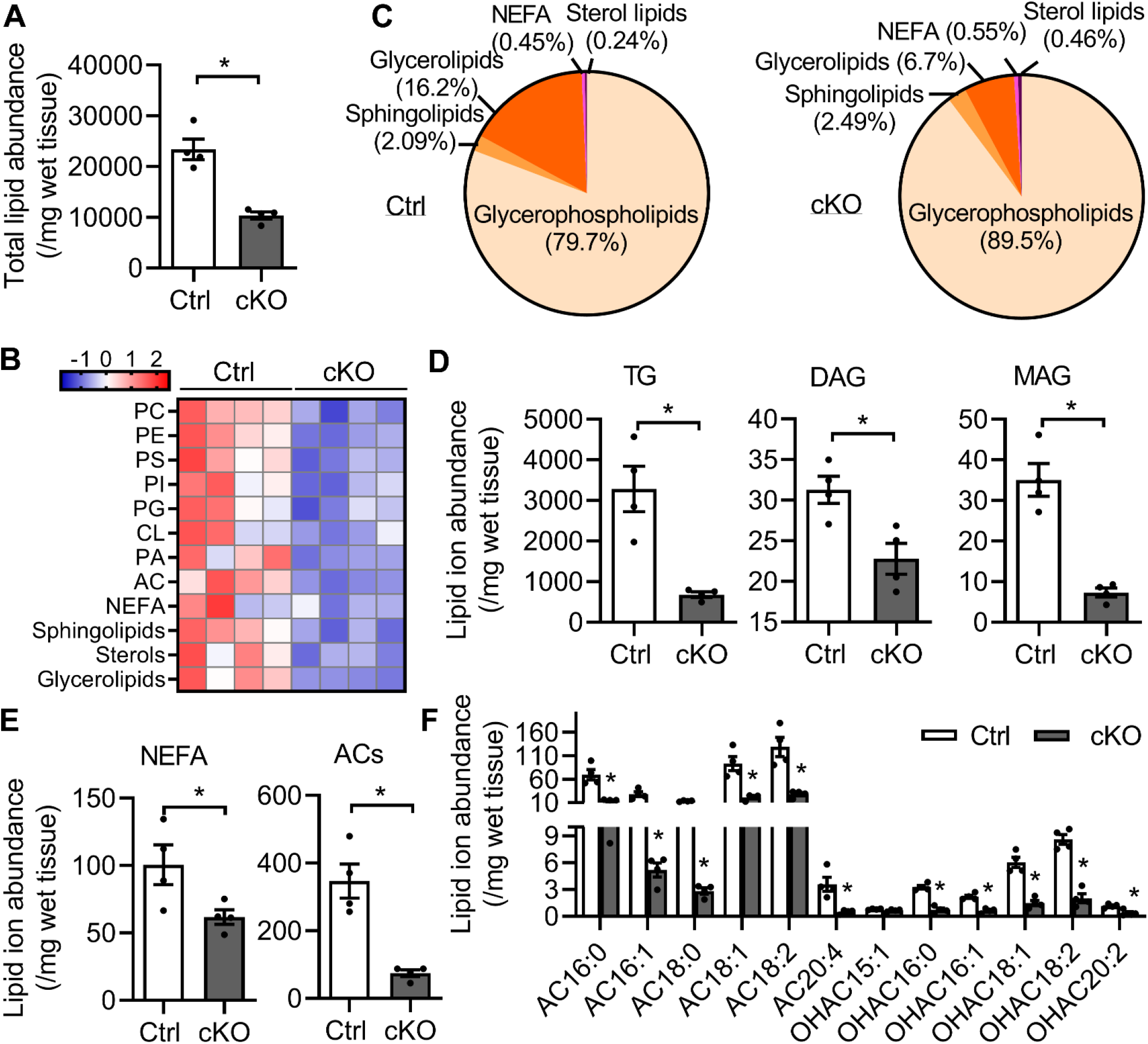
*Bscl2^cKO^* mice develop massive cardiac lipid remodeling and exhibit reduced metabolic reserve. **(A)** The total lipid ion abundance normalized to tissue weight. **(B)** Heatmap of lipid species including glycerophospholipids [i.e. phosphatidylcholine (PC), phosphatidylethanolamine (PE), phosphatidylserine (PS), phosphatidylinositol (PI), phosphatidylglycerol (PG), cardiolipin (CL), phosphatidic acid (PA), and acylcarnitines (AC)], nonesterified free fatty acids (NEFA), sphingolipids, sterols and glycerolipids based on Z-scores calculated from the summed ion abundances normalized to tissue weight. **(C)** Pie chart representing the proportional distribution of summed ion abundances of glycerolipid, glycerophospholipid, sphingolipid, NEFA, and sterol lipid classes. **(D-F)** Comparison of the total normalized ion abundances for **(D)** glycerolipids including TG, DAG and MAG, **(E)** NEFA and ACs, and **(F)** specific AC and hydroxyl acylcarnitines (OHAC) species. Global lipidomic analysis of ventricles by shotgun mass spectrometry was performed in nonfasting 6-month-old male *Cre-*; *Bscl2^f/f^* (Ctrl) and *Bscl2^cKO^* (cKO) mice. *n*=4 per group with each pooled from 3 animals. *: *P*< 0.05; **: *P*< 0.005, unpaired t tests (nonparametric).

To exclude the effect of the chronic expression of Cre transgene on massive lipid remodeling in *Bscl2^cKO^* mice, we included both Ctrl and *Cre+; Bscl2^w/w^* mice as controls to compare their cardiac TG contents and protein expression at 6 months of age. As expected, no differences were observed between Ctrl and *Cre+; Bscl2^w/w^* hearts, and only *Bscl2^cKO^* hearts displayed reduced TG (Supplemental Figure S3A) and increased ATGL expression (Supplemental Figure S3B). In addition, we observed similar expression of PPARα and its target proteins (CD36, CPT1β) between three genotypes (Supplemental Figure S3B). Collectively, these data emphasize a BSCL2-specific regulation of lipid remodeling in *Bscl2^cKO^* hearts independent of transcriptional activation of PPARα.

### Cardiomyopathy in 6-month-old *Bscl2^cKO^* mice is associated with energy deficiency independent of intrinsic mitochondrial dysfunction and oxidative stress

We next examined whether the massive lipid remodeling exerts maladaptive effects on mitochondrial function, leading to the development of HF in 6-month-old *Bscl2^cKO^* mice. Transmission electron microscopy images revealed a complete lack of LDs in 6-month-old *Bscl2^cKO^* hearts, in support of cardiac TG reduction (Figure 4A). Sarcomere arrangement, mitochondrial morphology and sizes, as well as mitochondria distribution were generally preserved in both Ctrl and *Bscl2^cKO^* hearts (Figure 4A). We performed respirational analysis of isolated mitochondria from hearts of Ctrl and *Bscl2^cKO^* mice in the presence of exogenous substrates (succinate), and identified similar basal and maximal oxygen consumption rates (Figure 4B). There were also no differences in coupled (ADP-driven) and uncoupled respiration between two genotypes, suggesting lack of obvious mitochondrial dysfunction (Figure 4C). In line with that, BSCL2 deletion did not affect the levels of marker proteins for mitochondrial ETC complexes expressed per μg total mitochondrial protein (Figure 4D). The absence of mitochondrial dysfunction and biogenesis was also confirmed in 6-month-old *Bscl2^cKO^* hearts as evidenced by similar expression of ETC complex proteins, PGC1α and mitochondrial stress marker Prohibitin compared with both Ctrl and *Cre+; Bscl2^w/w^* hearts (Supplemental Figure S4A). Despite an increased FAO in 3-month-old *Bscl2^cKO^* hearts, oxidation of 2’, 7’-dichlorofluorescein diacetate (DCFDA) was not augmented in heart extracts of 6-month-old *Bscl2^cKO^* mice (Supplemental Figure S4B). The level of lipid peroxides malondialdehyde (MDA) was even reduced in *Bscl2^cKO^* hearts compared with Ctrl hearts (Supplemental Figure S4C). Consistent with lack of oxidative stress, there were no differences in the expression of SOD1, SOD2 and Catalase in *Bscl2^cKO^* hearts relative to both Ctrl and *Cre+; Bscl2^w/w^* hearts (Supplemental Figure S4B). Together, these data suggest mitochondrial dysfunction and oxidative stress are unlikely to play a role in maladaptive cardiac remodeling and progression of HF in *Bscl2^cKO^* mice.

**Figure 4.**
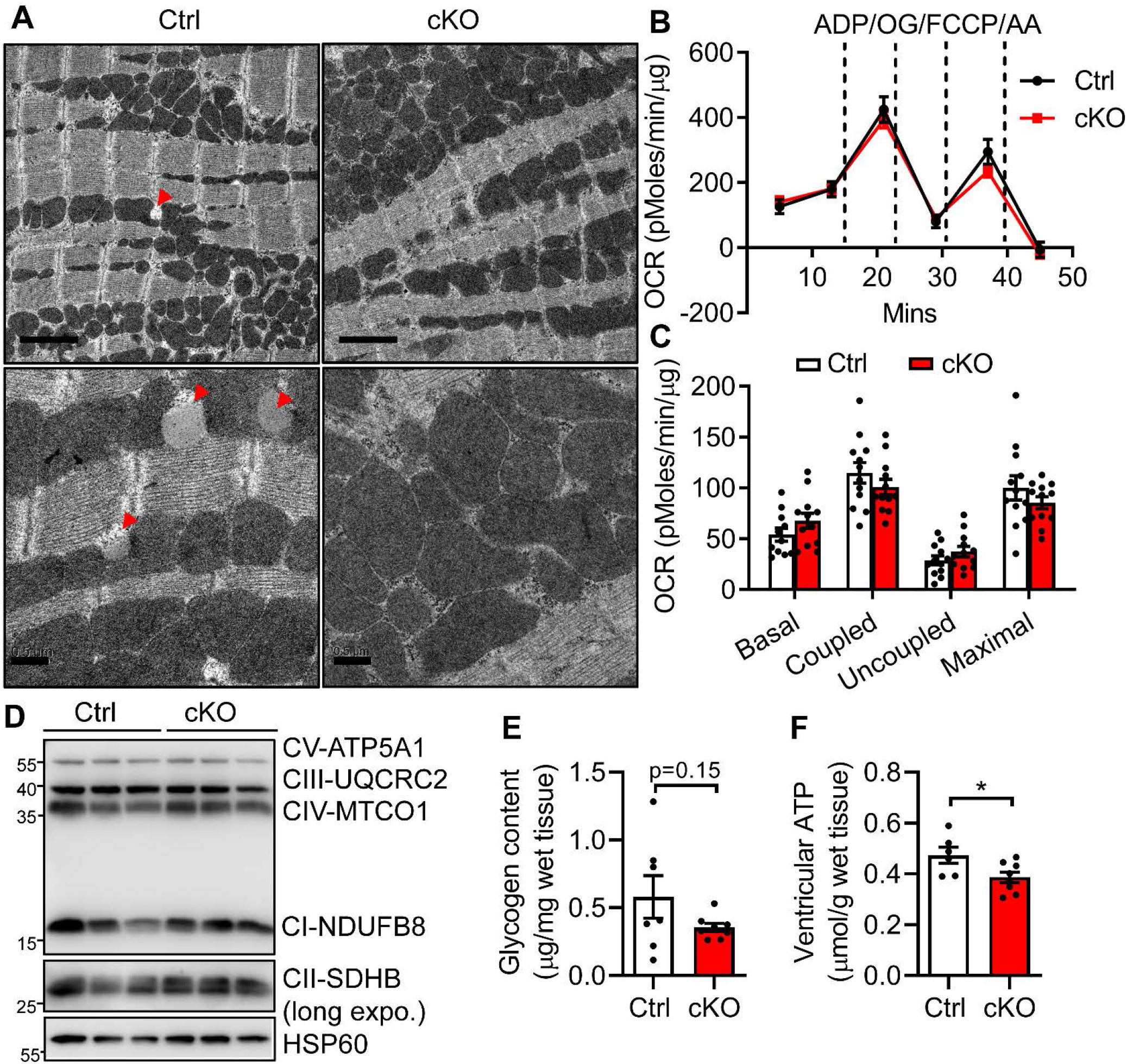
Cardiac dysfunction in *Bscl2^cKO^* mice is associated with bioenergetics failure independent of mitochondrial respiratory dysfunction. **(A)** Representative transmission electron micrographs of ventricles from 6-month-old *Cre-*; *Bscl2^f/f^* (Ctrl) and *Bscl2^cKO^* (cKO) mice. Red arrowheads: lipid droplets (LDs). Upper panels: scale bar = 2 μm; lower panels: scale bar = 0.5 μm. **(B-C)** Succinate/rotenone driven mitochondrial oxygen consumption rates (OCR) were measured by Seahorse XF24 analyzer with sequential addition of ADP, oligomycin (OG), FCCP and antimycin A (AA). Basal, coupled, uncoupled and maximal mitochondrial respiration were shown in **(C)**. *n*=12 per group. **(D)** Representative Western blotting in isolated mitochondria from ventricles of 6-month-old Ctrl and *Bscl2^cKO^* mice with mitochondrial HSP60 as controls. *n*=3 per group. Three independent experiments. **(E)** Ventricular glycogen contents, *n*=7 per group, and **(F)** ventricular ATP contents in nonfasting 6-month-old male *Ctrl* and *Bscl2^cKO^* mice. Ctrl, *n*=6; cKO, *n*=8. *: *P*< 0.05, unpaired t tests (parametric).

Interestingly, in contrast to 3-month-old *Bscl2^cKO^* mice (Figure 2J), we identified a tendency toward lower glycogen content in 6-month-old *Bscl2^cKO^* hearts (Figure 4E). There were no differences in the plasma concentrations of glucose and lipid substrates (TG, cholesterol, NEFA and glycerol) between 6 month-old Ctrl and *Bscl2^cKO^* mice (Supplemental Table S2). Despite the availability of extracellular substrates, the ventricular ATP content was significantly lower in hearts of 6-month-old *Bscl2^cKO^* mice (Figure 4F), in line with its reduced metabolic reserve (i.e. lipid and glycogen stores). Collectively, our data suggest chronic higher FAO caused by BSCL2 deletion in hearts leads to cardiac nutritional deprivation which is likely responsible for the cardiac energetic and contractile failure in 6-month-old *Bscl2^cKO^* mice.

### Inhibition of FAO reprograms cardiac lipidome and partially rescues cardiac dysfunction in *Bscl2^cKO^* mice

Since cardiac dysfunction in *Bscl2^cKO^* mice is associated with higher FAO, we treated mice with TMZ, a specific 3-Ketoacyl-CoA thiolase (3-KAT) inhibitor that inhibits FAO^33^, for up to 6 weeks starting at 6-month-old when *Bscl2^cKO^* mice already developed cardiac dysfunction. TMZ did not alter body weight (Supplemental Figure S5A) or liver weight (Supplemental Figure S5B), but tended to increase white fat mass in both genotypes (Supplemental Figure S5C). TMZ elevated circulating cholesterol levels in both Ctrl and *Bscl2^cKO^* mice, but there were no differences in plasma glucose, NEFA, glycerol or TG levels in the experimental groups (Supplemental Table S3). Ctrl and *Bscl2^cKO^* mice also demonstrated similar ventricle to body weight (Supplemental Figure S5D) and tibia length (Figure 5A) ratios after TMZ treatment. TMZ caused no changes in cardiac dilation and contractility of Ctrl mice, but was able to significantly improve wall thickness and cardiac contractility without notably normalizing dilatation of *Bscl2^cKO^* hearts (Figure 5B-D and Supplemental Table S3). Attenuation of cardiac dysfunction by TMZ was also evident in female *Bscl2^cKO^* mice (Supplemental Figure S5E-H).

**Figure 5.**
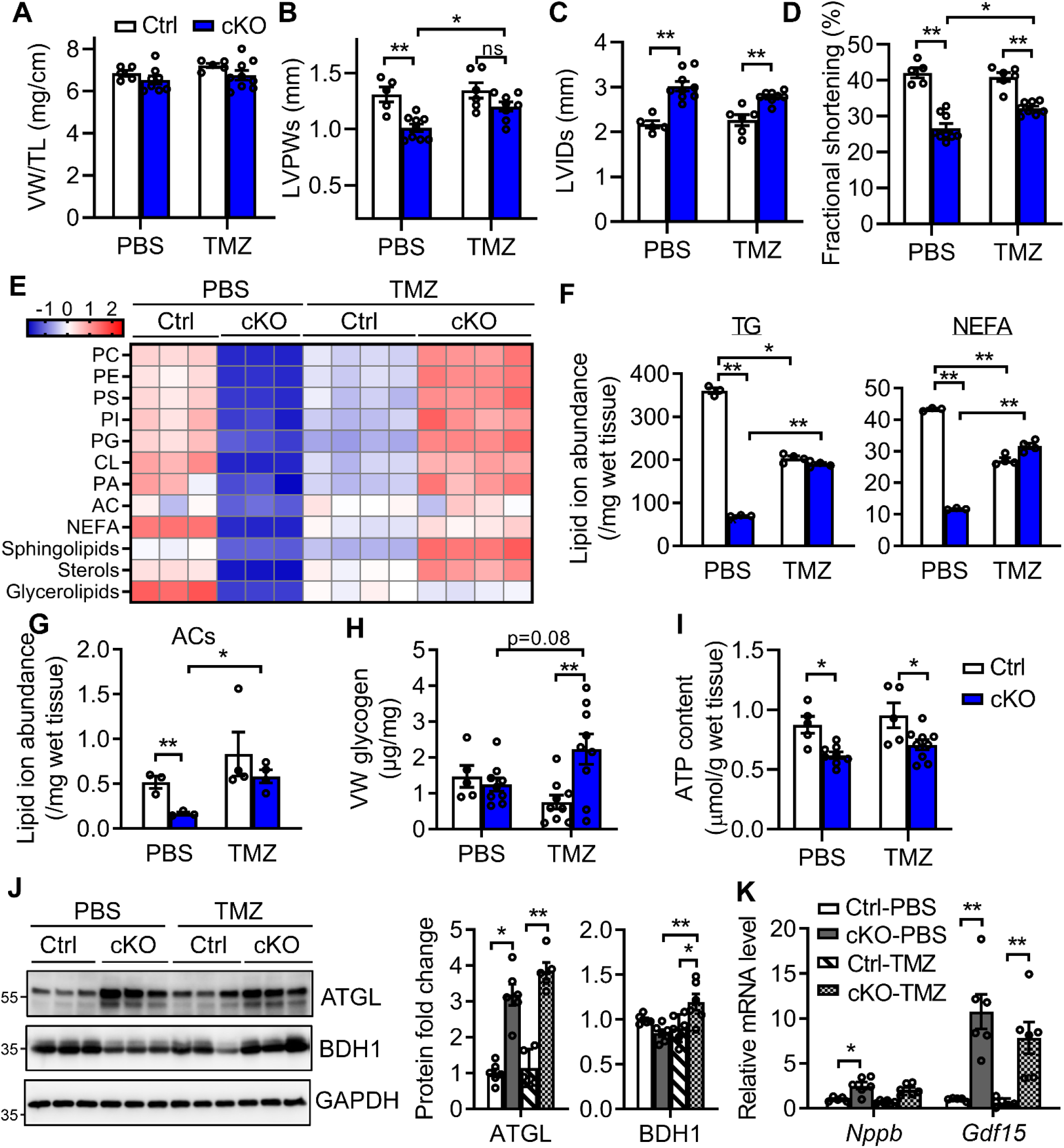
Inhibition of fatty acid oxidation partially improves metabolic reserve and cardiac function in *Bscl2^cKO^* mice. 6-month-old male *Cre-*; *Bscl2^f/f^* (Ctrl) and *Bscl2^cKO^* (cKO) mice received daily i.p. injection with PBS or trimetazidine (TMZ) at 15 mg/kg body weights (BW) for a total of 6 weeks. **(A)** Ratio of ventricle weight (VW) to tibia length (TL). Ctrl, *n*=5; cKO, *n*=9. **(B)** Left ventricle post wall thickness at end systole (LVPWs, mm); **(C)** left ventricular internal diameter in end systole (LVIDs); and **(D)** fractional shortening. Ctrl, *n*=6; cKO, *n*=9. **(E)** Heatmap of major lipid species based on Z-score calculated from the summed ion abundances normalized to milligram of wet tissue; **(F-G)** comparison of the total normalized ion abundances for **(F)** glycerolipids and NEFA as well as **(G)** ACs. *n*=3 per PBS-treated group. *n*=4 per TMZ-treated group. Each sample was pooled from 3 animals. **(H)** Ventricular glycogen. PBS-Ctrl, *n*=5; PBS-cKO, *n*=9. *n*=9 per TMZ-treated group. **(I)** Ventricular ATP contents normalized to gram of wet tissue. PBS-Ctrl, *n*=5; PBS-cKO, *n*=8. TMZ-Ctrl, *n*=5, TMZ-cKO, *n*=9. **(J-K)** Representative Western blotting and quantification of total cell extracts from ventricles of 6-month-old PBS and TMZ-treated Ctrl and *Bscl2^cKO^* mice. *n*=6 per group. Data were normalized to GAPDH with PBS-treated Ctrl set to 1. Two independent experiments. **(L)** RT-PCR analysis of cardiac stress genes in ventricles of 6-month-old PBS and TMZ-treated male mice. *n*=6 per group. *: *P*< 0.05; **: *P*< 0.005. Two-way ANOVA followed by Tukey’s post-hoc tests.

We performed another independent set of untargeted lipidomics to examine the effect of TMZ on cardiac lipidome. As indicated in Figure 5E, PBS-treated *Bscl2^cKO^* hearts recapitulated all changes in cardiac lipid contents as we previously observed in 6-month-old *Bscl2^cKO^* hearts (Figure 3). TMZ unexpectedly reduced the absolute levels of cardiac lipid contents in all categories except ACs in Ctrl hearts. In contrast, TMZ dramatically enhanced the accumulation of these lipid classes in *Bscl2^cKO^* hearts relative to PBS-treated *Bscl2^cKO^* hearts. Especially, the absolute levels of phospholipids, sphingolipids, and sterols were even greater in TMZ-treated *Bscl2^cKO^* than PBS-treated Ctrl hearts (Figure 5E and Supplemental Excel 2). Specifically, TMZ was able to markedly increase TG, NEFAs and ACs in *Bscl2^cKO^* hearts (Figure 5F-G). Moreover, TMZ slightly lowered glycogen in Ctrl hearts. However, it caused more glycogen accumulation in *Bscl2^cKO^* hearts (Figure 5H). Despite a significant upregulation of metabolic reserve, there was only a very minimal but non-significant improvement of ventricular ATP content in TMZ-treated *Bscl2^cKO^* hearts (Figure 5I). As expected, TMZ itself exerted no effect on ATGL upregulation in *Bscl2^cKO^* hearts, since it acts on FAO downstream of ATGL (Figure 5J). However, the expression of BDH1, a crucial enzyme responsible for cardiac ketolysis, was significantly higher in *Bscl2^cKO^* hearts after TMZ treatment (Figure 5J). There was also a trend for TMZ to attenuate the gene expression of cardiac stress markers in *Bscl2^cKO^* hearts (Figure 5K). Together, our data suggest inhibiting FAO in *Bscl2^cKO^* hearts is able to partially restore cardiac function, potentially through modifying cardiac metabolism.

### HFD significantly improves cardiac energy substrates and prevents heart failure in *Bscl2^cKO^* mice

To gain further insight into the importance of metabolic reserve in energy-starved *Bscl2^cKO^* hearts, we also fed 3-month-old Ctrl and *Bscl2^cKO^* male mice a normal chow diet (NCD) or a 60% high fat diet (HFD) for a period of 3 months. By 6 months old, Ctrl and *Bscl2^cKO^* mice exhibited similar obese phenotype with comparable weight gain after HFD feeding (Supplemental Table S2). They exhibited similarly higher levels of plasma glucose, cholesterol, NEFA and glycerol relative to their NCD-fed counterparts, and there were no differences in serum TG concentrations in all four groups (Supplemental Table S2). HFD slightly increased the ventricle weight to tibia length ratios in both genotypes, indicating mild but comparable cardiac hypertrophy (Figure 6A). We found no significant changes in wall thickness, dilation and contractility in Ctrl mice after 3 months of HFD feeding (Figure 6B-D and Supplemental Table S2). However, HFD was able to increase LVPWs and LVAWs, and improve cardiac contractility of *Bscl2^cKO^* mice to the similar levels of HFD-fed Ctrl mice despite exerting no effect on dilation (Figure 6B-D and Supplemental Table S2).

**Figure 6.**
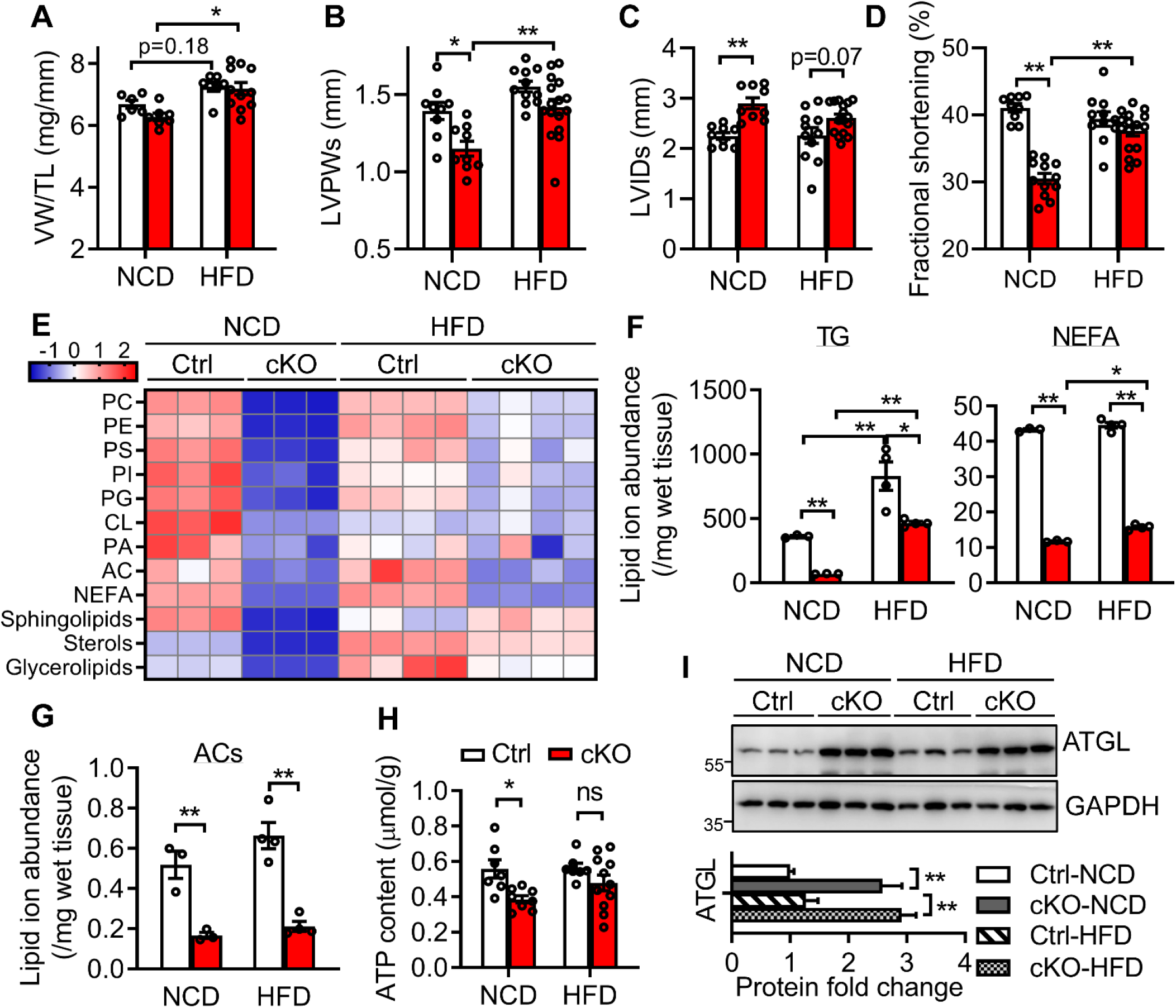
High fat diet feeding increases metabolic reserve and prevents cardiac dysfunction in *Bscl2^cKO^* mice. 3-month-old *Bscl2^f/f^* (Ctrl) and *Bscl2^cKO^* (cKO) mice were fed with normal chow diet (NCD) or high fat diet (HFD, 60% fat calories) for a total of 3 months. **(A)** ratio of ventricle weight (VW) to tibial length (TL). Ctrl, *n*=5; cKO, *n*=9. **(B)** Left ventricle post wall thickness at end systole (LVPWs, mm); **(C)** left ventricular internal diameter in end systole (LVIDs); and **(D)** fractional shortening. NCD-Ctrl, *n*=9; NCD-cKO, *n*=12; HFD-Ctrl, *n*=12; HFD-cKO, *n*=16. **(E)** Heatmap of major lipid species based on Z-score calculated from summed ion abundances normalized to milligram of wet tissue; **(F-G)** comparison of the total normalized ion abundances for **(F)** glycerolipids and NEFA as well as **(G)** ACs. *n*=3 per NCD group. *n*=4 per HFD group. Each sample was pooled from 3 animals. **(H)** Myocardial ATP content as normalized to gram of wet tissue. NCD-Ctrl, *n*=8; NCD-cKO, *n*=8; HFD-Ctrl, *n*=7; HFD-cKO, *n*=11. **(I)** Representative Western blotting of total cell extracts from ventricles of Ctrl and *Bscl2^cKO^* mice. *n*=3 per group. Two independent experiments. *: *P*< 0.05; **: *P*< 0.005. Two-way ANOVA followed by Tukey’s post-hoc tests.

We also determined cardiac lipidome of HFD-fed hearts with the above PBS-treated hearts as NCD controls. Interestingly, in Ctrl hearts, we found HFD reduced the levels of phospholipids (i.e. PC, PS, PI, PG, CL and PA) and sphingolipids, but preferentially increased the levels of sterols and glycerolipids when comparing with NCD (Figure 6E). The amounts of almost all lipid classes were increased in *Bscl2^cKO^* hearts after HFD feeding, albeit still lower than HFD-fed Ctrl hearts (Figure 6E and Supplemental Excel 2). Specifically, TG levels were greatly upregulated in both Ctrl and *Bscl2^cKO^* hearts after HFD feeding; whereas the levels of NEFA and ACs in *Bscl2^cKO^* hearts were increased to a lesser extent by HFD when compared with NCD (Figure 6F-G). Nevertheless, while *Bscl2^cKO^* hearts contained less ATP under NCD, the ventricular ATP contents in HFD-fed *Bscl2^cKO^* mice was almost restored to the levels of HFD-fed Ctrl hearts, suggesting improved cardiac energetics (Figure 6H). Immunoblot analysis revealed similar upregulation of ATGL in *Bscl2^cKO^* relative to Ctrl hearts regardless of diets (Figure 6I). Collectively, HFD could restore cardiac function by improving cardiac energetics via increasing energy supply in *Bscl2^cKO^* mice.

## Discussion

In this study, we show that genetic deletion of BSCL2 in cardiomyocytes leads to dramatic cardiac lipid remodeling and dilated HF in mice. Mechanistically, cardiac BSCL2 ablation causes ATGL overexpression, excessive FAO and overt cardiac lipid remodeling. Increased lipid catabolism ultimately exhausts intramyocellular lipid and glycogen reserve and is likely responsible for the energetic and contractile failure in *Bscl2^cKO^* mice. Importantly, inhibiting FAO by promoting substrate switch or HFD feeding via increasing lipid supply alleviate cardiac dysfunction in *Bscl2^cKO^* mice. Our results thus identify a novel and indispensable role of BSCL2 in regulating a preferential oxidation of FAs from endogenous cardiac TG lipolysis which governs cardiac energetics and function.

BSCL2 deletion enhances cAMP/PKA triggered ATGL-mediated lipolysis and FAO in adipose tissue^30–32^. Similar to global *Bscl2^−/−^* hearts^27^, *Bscl2^cKO^* hearts exhibited higher ATGL expression associated with reduced TG contents and accelerated FAO, highlighting the cell-autonomous role of cardiac BSCL2 in lipid catabolism despite its low expression in hearts. Interestingly, *Bscl2^cKO^* mice developed more severe cardiomyopathy than global *Bscl2^−/−^* mice, suggesting differential pathological mechanisms. Lipodystrophic *Bscl2^−/−^* mice undergo mild hypertrophic cardiomyopathy, which can be attenuated by partially restoring fat mass and/or improving whole-body and cardiac insulin resistance^27, 34^. This suggests insulin resistance largely accounts for the pathophysiology of metabolic cardiomyopathy in lipodystrophic *Bscl2^−/−^* mice, resembling diabetic hearts. On the other hand, *Bscl2^cKO^* mice developed energy deficit-induced dilated HF which was independent of profound remodeling of structural changes (e.g. hypertrophy and fibrosis) (Figure 1) and insulin resistance (data not shown). We speculate that hyperphagia in lipodystrophic *Bscl2^−/−^* mice may actually provide more circulating lipid metabolites which protect BSCL2-deleted heart from cardiac substrate exhaustion and HF as we observed in *Bscl2^cKO^* mice. Notably, a previous report identified no differences in cardiac function of 55-60 week-old mice with cardiac deletion of BSCL2 driven by the same Myh6-Cre^35^. This discrepancy could be due to differences in construct design or background strain.

Comparisons of cardiac lipidomes between *Bscl2^−/−^* and *Bscl2^cKO^* mice clearly demarcate the autonomous and non-autonomous effects of BSCL2 deletion on lipid remodeling. *Bscl2^cKO^* mice exhibited an almost 50% reduction in total lipid contents concomitant with downregulation of a wide array of lipid species from glycerolipids to phospholipids to ACs (Figure 3). On the contrary, *Bscl2^−/−^* hearts only demonstrated reduced glycerolipid contents with a tendency toward lower levels of glycerophospholipids, NEFA and ACs ^27^. Such discrepancy could again be attributed to the excess circulating FA levels in hyperphagic *Bscl2^−/−^* mice. Notably, the levels of cardiac PA, the important glycerolipid intermediates and phospholipid precursors, were consistently reduced in both *Bscl2^−/−^* and *Bscl2^cKO^* hearts, suggesting a potential specific role of BSCL2 in regulating PA metabolism. Different from our results, PA levels were shown to be enriched in BSCL2-deleted murine adipose tissue^36^ and yeasts^37, 38^. Whether this can be ascribed to differences in tissue depots or species is not known. Additionally, BSCl2-deleted *Drosophila* S2 cells also displayed reduced phospholipids which potentially account for the formation of giant lipid droplets^39^. More work is needed to understand whether BSCL2 plays a direct role in mediating phospholipid metabolism.

PGC-1α and PPARα are key activators of TG dynamics and content in the heart^40, 41^. Especially, ATGL-mediated fat catabolism has been directly linked to cardiac PGC-1 and PPARα expression as well as FAO rates^8^. While *Bscl2^cKO^* hearts clearly exhibited excessive FAO in *ex vivo* perfused working hearts preceding impaired cardiac function, none of PGC-1α and PPARα and their downstream target genes were altered (Figure 2). Interestingly, lean mice with myocardial ATGL overexpression^10, 11^ or acetyl CoA carboxylase 2 (ACC2) deletion^42^ maintain normal cardiac energetics and performance despite higher TG turnover or FAO rates. Thus, it needs to be recognized that other changes in cellular metabolism independent of ATGL-mediated intramyocardial lipid catabolism may exist to contribute to the cardiac energy deficit and progressive deterioration of cardiac function in *Bscl2^cKO^* mice at baseline, which warrants further investigation. Regarding the posttranscriptional regulation of ATGL, ATGL is known to be ubiquitinated [^43^ and data not shown], and we have previously demonstrated enhanced ATGL stability in *Bscl2^−/−^* cardiomyocytes and mouse embryonic fibroblasts (MEFs)^27^. However, we were not able to pull down endogenous ATGL using current available antibodies, which prevents us from examining ATGL ubiquitination in BSCL2-deleted hearts. Therefore, the molecular events for the posttranscriptional regulation of ATGL in the absence of BSCL2 remain to be identified.

Reliance on FAO in obesity and/or diabetes is correlated with lower cardiac efficiency, impaired mitochondrial respiratory function and increased ROS production^44^. Alteration of the cardiac lipidome may also mediate functional impairment through dampening mitochondrial function^45^. However, none of these abnormalities occurred in our *Bscl2^cKO^* mice despite massive reduction of cardiac lipid contents. In fact, BSCL2-deleted mitochondria maintain efficient oxidative phosphorylation in the presence of exogenous substrates (Figure 4). This implies that the ATP deficit in *Bscl2^cKO^* hearts is mainly due to insufficient endogenous substrate *in vivo* independent of the intrinsic defects in mitochondrial function. Higher cardiac FAO is normally associated with increased exogenous lipid import as observed in diabetic hearts or hearts with cardiac-restricted overexpression of PPARα^46^. However, our *Bscl2^cKO^* mice displayed similar expression of lipid uptake genes and comparable circulating lipid metabolites, suggesting no defect in lipid transport. This may ultimately result in an imbalance of lipid consumption and supply within *Bscl2^cKO^* myocardium leading to downregulation of vast amounts of lipid substrates (Figure 3). In addition, AC levels tightly reflect the FAO rates, and AC profiling has been used to identify FAO dysregulation^47^. Previous reports suggest muscle AC levels correlate negatively with FAO in the postabsorptive state^48^. Notably, *Bscl2^cKO^* hearts demonstrated no alterations in CPT1 expression, suggesting intact carnitine shuttle (Figure 3K). The almost unanimous reduction of long-chain ACs in *Bscl2^cKO^* hearts highlights the presence of increased FAO which may eventually deplete mitochondrial ACs thus reducing substrates entering TCA cycles and causing energy deficit. Thus, our *Bscl2^cKO^* mice constitute as the first animal model that demonstrates excessive myocardial lipid catabolism associated with progressive deterioration of metabolic reserve and HF in the absence of elevated lipid uptake.

TMZ has been reported to improve cardiac function in experimental models of ischemia/reperfusion injury^49–51^ and ischemic HF patients^52–54^. It predominantly acts by shifting energy production from NEFA to glucose oxidation in the heart^50^. TMZ exerts cardioprotective role in our *Bscl2^cKO^* mice. Yet, the underlying mechanisms may not be simply explained by alleviation of energy deficit, as TMZ failed to significantly improve intracellular ATP levels in *Bscl2^cKO^* hearts. Although our focus was primarily on TMZ effect on cardiac function, we surprisingly uncovered that TMZ remodels cardiac lipidome by downregulating the lipid contents of almost all lipid classes in normal mouse hearts. We speculate inhibiting FAO may also put a brake on lipid transport and/or synthesis in the heart. Conversely, TMZ promotes drastic lipid and glycogen accumulation in metabolic-stressed *Bscl2^cKO^* hearts. The prominence of glycogen in TMZ-treated *Bscl2^cKO^* hearts may reflect enhanced glucose utilization as glycogen content has also been shown to commensurate with augmented carbohydrate metabolism such as in GLUT1-overexpressing hearts^55^. Notably, none of these changes can be associated with differences in the protein expression of lipid and glucose metabolic genes (data not shown). Interestingly, aside from glucose utilization, TMZ may also trigger enhanced cardiac ketolysis in *Bscl2^cKO^* hearts which demonstrated higher BDH1 expression. This finding concurs with a recent report on TMZ’s induction of cardiac β-hydroxybutyrate flux to attenuate isopropanol-induced rat heart failure^56^. Whether TMZ indeed induces substrate switch to both glucose and ketone bodies in *Bscl2^cKO^* hearts needs to be further dissected through *ex vivo* perfused working hearts. In addition, we cannot exclude the nonmetabolic effects of TMZ in preventing left ventricle remodeling independent of its inhibitory activity on FAO^57, 58^. Nevertheless, our study underscores that TMZ has potential in ameliorating cardiac function and slows HF progression in a non-ischemic model of HF.

HFD alone has been shown to be cardioprotective especially in alleviating energy-compromised HF^59^. In the present study, we find that decreased cardiac function in *Bscl2^cKO^* mice can be attenuated by HFD feeding. Lipidomics study further confirmed improved cardiac metabolic substrates mainly in the form of TG in HFD-fed *Bscl2^cKO^* mice. This was in line with the notion that HFD feeding provides more coronary circulation of substrates to match up the rate of enhanced lipid utilization thus attenuating the myocardial ATP deficit in the energy-deprived failing *Bscl2^cKO^* hearts. In spite of an improvement in cardiac contractility, we were not able to observe a significant reduction of cardiac stress markers in HFD-treated *Bscl2^cKO^* mice (not shown). Nonetheless, results from our HFD feeding studies support the cardio-protective effect of HFD on the energy-deprived HF.

In conclusion, our study highlights an important link between BSCL2 and myocardial energy metabolism and function and advances our understanding of the relationship between TG dynamics, FAO rates and the pathogenesis of HF. It may also provide insights into the therapeutic approaches in the treatment of cardiac disorders related to dysregulated metabolism in general.

## Materials and Methods

### Mice

*Cre+; Bscl2^f/f^* mice (designated as *Bscl2^cKO^*) mice were generated by breeding *Bscl2^f/f^* mice^13^ with transgenic mice expressing Cre recombinase under the cardiac-specific alpha myosin-heavy chain 6 (*Myh6*) promoter (JAX Cat#: 011038)^60^. The *Myh6-Cre* mice (designated as *Cre+; Bscl2^w/w^*) were included as controls when assessing cardiac functions. *Cre-; Bscl2^f/f^* mice (designated as Ctrl) were used as controls for the majority of the studies. To inhibit mitochondrial β-oxidation, 6-month-old Ctrl and *Bscl2^cKO^* mice were i.p. injected with PBS or trimetazidine (TMZ) at 15 mg/kg/day for 6 weeks, a dose that does not induce whole-body insulin resistance^33^. Ctrl and *Bscl2^cKO^* mice were fed a 60% HFD (Research Diets; D12492) starting at 3 months of age for a duration of 3 months. Most experiments were performed in *ad libitum* male mice and repeated in female mice. All mice were housed in the central animal facility with room temperature controlled at 21°C, and an artificial 12 h:12 h light: dark cycle (lights on at 06:00 am). Mice were directly sacrificed by cervical dislocation and hearts were rapidly excised. In ex vivo perfused working heart experiments, hearts were rapidly excised (<30 s) following anesthesia via intraperitoneal ketamine/xylazine (80/10 mg/kg) injection. All procedures involving animals and tissues conform to the NIH Guide for the Care and Use of Laboratory Animals and were approved by the Institutional Animal Care and Use Committee at the Augusta University.

### Echocardiography

Echocardiographic analyses were performed on mice under anesthesia which was first induced with 5% isoflurane in 100% O_2_ for 1 min in a vaporizer then maintained during spontaneous breathing of 1.25% isoflurane in 100% O_2_ at a flow rate of 1 l/min via a small nose cone. Two-dimensional guided M-mode echoes were obtained from short-and long-axis views at the levels of the largest left ventricular diameter using a a VisualSonics Vevo 2100 echocardiography machine equipped with a 30 MHz probe (VisualSonics). All measurements and calculations were done in triplicates.

### Tissue TG analyses

For tissue TG enzymatic analyses, lipids were extracted from tissue homogenates and dissolved in chloroform. The concentrations of TG were measured using a triglyceride assay kit (Infinity™ triglycerides kit, Thermo Fisher Scientific) and normalized to tissue weights as previously described^13^.

### Histology and transmission electron microscopy (TEM)

Whole hearts were fixed, embedded and cut along the coronal plane to visualize the four-chamber view. The apex of left ventricle (LV) tissue was fixed and stained for electron microscopical imaging in a JEM 1230 transmission electron microscope (JEOL USA Inc., Peabody, MA) at 110 kV with an UltraScan 4000 CCD camera & First Light Digital Camera Controller (Gatan Inc., Pleasanton, CA) as previously described^27^.

### Lipidomic analysis by high resolution/accurate mass spectrometry and tandem mass spectrometry

Total lipids from frozen ventricles were extracted and resuspended in isopropanol:methanol:chloroform (4:2:1 v:v:v) containing 20 mM ammonium formate followed by untargeted lipidomic analysis. Relative quantification of abundances between samples was performed by normalizing target lipid ion peak areas to the PC (14:0/14:0) internal standard followed by normalization to tissue weights as previously described^27^.

### FAO assays

FAO reaction assays with LV homogenates were prepared and carried out as detailed previously ^61^. Briefly, ≈ 25 mg pieces of freshly isolated ventricular tissues were placed in STE buffer (250 mM Sucrose, 10 mM Tris, pH=7.5 and 1 mM EDTA) and homogenized using a glass dounce homogenizer (20 loose and 20 tight strokes). 20 μL tissue homogenates were incubated with 380 μL reaction buffer containing 1 μCi/mL [1–^14^C] palmitic acid substrate for fatty acid oxidation. The released [^14^C]CO_2_ was captured by hydroamine soaked filter paper and measured by scintillation counting, while acid soluble metabolites (ASM) were analyzed by centrifugation and counting of ^14^C radioactivity in the supernatant. Data were normalized to the total protein contents for LV homogenates.

### Isolation and culture of adult cardiomyocytes

The isolation of adult mouse cardiomyocytes was carried out based on established procedures ^62^. The cardiomyocytes were suspended in plating media and plated onto laminin (5 μg/mL) precoated tissue culture plates. 1 h after plating, myocytes were changed to the culture media in the absence of 2,3-butanedione monoxime (BDM), ITS (Insulin/transferrin/selenium supplement) and lipid and kept in culture for 4 h, before exposure to isoproterenol (1 μM) for 20 mins.

### Substrate metabolism in isolated working mouse hearts

Myocardial substrate utilization and contractile function were measured *ex vivo* in hearts isolated from 14 to 16 week-old male Ctrl and *Bscl2^cKO^* mice. All hearts were prepared and perfused in the working mode (non-recirculating manner) for 30 minutes with a preload of 12.5 mmHg and an afterload of 50 mmHg as previously described^63, 64^. Standard Krebs–Henseleit buffer was supplemented with 8 mM glucose, 0.4 mM oleate conjugated to 3%BSA (fraction V, FA-free; dialyzed), 10 μU/ml insulin (basal/fasting concentration), 0.05 mM L-carnitine, and 0.13 mM glycerol. Metabolic fluxes were assessed through the use of distinct radiolabeled tracers: 1) [U-^14^C]-glucose (0.12 mCi/L from MP Biomedicals; glucose oxidation); and 2) [9, 10-^3^H]-oleate (0.067 mCi/L from Sigma-Aldrich; β-oxidation). Measures of cardiac metabolism (*e.g*., oleate and glucose oxidation, and oxygen consumption) and function (*e.g*., cardiac power) were determined. At the end of the perfusion period, hearts were snap-frozen in liquid nitrogen and stored at −80°C prior to analysis. Data were presented as steady state values (*i.e*., the mean of the last two time points during a distinct perfusion condition for each individual heart).

### Mitochondrial isolation and measurement of mitochondrial respiration

Fresh ventricles were isolated and minced for mitochondrial isolation and measurement of mitochondrial respiration by XF24 Extracellular Flux Analyzer (Seahorse Bioscience) as previously described^32^. See Supplemental Materials and Methods for details.

### Tissue ATP measurement

ATP content was determined by using ATP Bioluminescent Assay Kit (FL-AA; Sigma-Aldrich, Saint Louis, MO, USA) according to the manufacturer’s procedure. Briefly, frozen heart tissues were homogenized in cold 10% trichloroacetic acid buffer, centrifuged at 5000 × g for 10 min at 4°C followed by neutralization with 50 mM Tris-acetate (pH 7.8). The ATP content was then determined by a multi-mode microplate reader with luminescence luminometer (FLUOstar Omega; BMG Labtech). Data were normalized to tissue weight.

### RNA isolation and real-time quantitative PCR

Total RNA was extracted with Trizol Reagent (Thermo Fisher) and reverse-transcribed using MLV-V reverse transcriptase using random primers (Invitrogen). Real-time quantitative RT-PCR was performed on the Strategene MX3005 system. Data were normalized to 2 housekeeping genes (*Actb* and *36B4*) based on Genorm algorithm (medgen.ugent.be/genorm/) and expressed as fold changes. All tissue gene expression studies were performed in nonfasted mice. RT-PCR Primers were listed in Supplemental Table S4.

### Immunoblotting

Tissues were lysed in lysis buffer containing 25 mM Tris-HCl (pH 7.4), 150 mM NaCl, 2 mM EDTA, 1% Triton X-100 and 10% glycerol with freshly added protease and phosphatase inhibitor cocktail (Sigma). The protein concentration was determined by Bradford protein assay (Bio-Rad). Western blotting and quantification were performed as previously described^27^. Specific antibodies were listed in Supplemental materials.

### Statistical analysis

Quantitative data were presented as means ± SEM. Animal experiments were performed with at least three independent cohorts. Statistical comparisons were made by using unpaired *t* test, one-way ANOVA followed by Dunnett’s multiple comparisons test, two-way ANOVA followed by Tukey’s post-hoc tests, or multiple *t* tests after correction using the Holm-Sidak method using the built-in statistics of GraphPad Prism 9 software. A *P* value of less than 0.05 was considered statistically significant.

Additional methodological details are included in the Supplemental Materials.

## Acknowledgement

We thank the Electron Microscopy and Histology Core at Augusta University for technical assistance and electron microscope imaging. This work was supported by National Heart, Lung and Blood Institute [1R01HL132182-01 to W.C.], National Institute of General Medical Sciences [R01GM124108 to J.L.], National Institute on Aging [R01AG049835 to J.L], and the American Heart Association Career Development Award (18CDA34080244 to H.Z.).

## Author contributions

H. Zhou and W. Chen conceived the project and designed the research and wrote the manuscript. H. Zhou performed most physiological, biochemical and molecular studies; Ji Li and M.E. Young performed and supervised ex vivo perfused working hearts. Todd A. Lydic performed lipidomic analysis and assisted with data interpretation. Jie Li and H. Su performed and supervised adult cardiomyocyte isolation. H. Su, Ji Li and M.E. Young helped editing the manuscript.

## References

1. Writing Committee M, Yancy CW, Jessup M, Bozkurt B, Butler J, Casey DE, Jr., Drazner MH, Fonarow GC, Geraci SA, Horwich T, Januzzi JL, Johnson MR, Kasper EK, Levy WC, Masoudi FA, McBride PE, McMurray JJ, Mitchell JE, Peterson PN, Riegel B, Sam F, Stevenson LW, Tang WH, Tsai EJ, Wilkoff BL, American College of Cardiology Foundation/American Heart Association Task Force on Practice G. 2013 ACCF/AHA guideline for the management of heart failure: a report of the American College of Cardiology Foundation/American Heart Association Task Force on practice guidelines. Circulation 2013; 128:e240–327.

2. Grynberg A, Demaison L. Fatty acid oxidation in the heart. J Cardiovasc Pharmacol 1996;28 Suppl 1:S11–17.

3. van der Vusse GJ, Glatz JF, Stam HC, Reneman RS. Fatty acid homeostasis in the normoxic and ischemic heart. Physiol Rev 1992;72:881–940.

4. Kolwicz SC, Jr., Tian R. Glucose metabolism and cardiac hypertrophy. Cardiovasc Res 2011;90:194–201.

5. Ingwall JS, Weiss RG. Is the failing heart energy starved? On using chemical energy to support cardiac function. Circ Res 2004;95:135–145.

6. Neubauer S, Horn M, Cramer M, Harre K, Newell JB, Peters W, Pabst T, Ertl G, Hahn D, Ingwall JS, Kochsiek K. Myocardial phosphocreatine-to-ATP ratio is a predictor of mortality in patients with dilated cardiomyopathy. Circulation 1997;96:2190–2196.

7. Goldberg IJ, Trent CM, Schulze PC. Lipid metabolism and toxicity in the heart. Cell Metab 2012;15:805–812.

8. Haemmerle G, Moustafa T, Woelkart G, Buttner S, Schmidt A, van de Weijer T, Hesselink M, Jaeger D, Kienesberger PC, Zierler K, Schreiber R, Eichmann T, Kolb D, Kotzbeck P, Schweiger M, Kumari M, Eder S, Schoiswohl G, Wongsiriroj N, Pollak NM, Radner FP, Preiss-Landl K, Kolbe T, Rulicke T, Pieske B, Trauner M, Lass A, Zimmermann R, Hoefler G, Cinti S, Kershaw EE, Schrauwen P, Madeo F, Mayer B, Zechner R. ATGL-mediated fat catabolism regulates cardiac mitochondrial function via PPAR-alpha and PGC-1. Nat Med 2011;17:1076–1085.

9. Zimmermann R, Strauss JG, Haemmerle G, Schoiswohl G, Birner-Gruenberger R, Riederer M, Lass A, Neuberger G, Eisenhaber F, Hermetter A, Zechner R. Fat mobilization in adipose tissue is promoted by adipose triglyceride lipase. Science 2004;306:1383–1386.

10. Pulinilkunnil T, Kienesberger PC, Nagendran J, Sharma N, Young ME, Dyck JR. Cardiac-specific adipose triglyceride lipase overexpression protects from cardiac steatosis and dilated cardiomyopathy following diet-induced obesity. Int J Obes (Lond) 2014;38:205–215.

11. Pulinilkunnil T, Kienesberger PC, Nagendran J, Waller TJ, Young ME, Kershaw EE, Korbutt G, Haemmerle G, Zechner R, Dyck JR. Myocardial adipose triglyceride lipase overexpression protects diabetic mice from the development of lipotoxic cardiomyopathy. Diabetes 2013;62:1464–1477.

12. Windpassinger C, Auer-Grumbach M, Irobi J, Patel H, Petek E, Horl G, Malli R, Reed JA, Dierick I, Verpoorten N, Warner TT, Proukakis C, Van den Bergh P, Verellen C, Van Maldergem L, Merlini L, De Jonghe P, Timmerman V, Crosby AH, Wagner K. Heterozygous missense mutations in BSCL2 are associated with distal hereditary motor neuropathy and Silver syndrome. Nat Genet 2004;36:271–276.

13. Chen W, Chang B, Saha P, Hartig SM, Li L, Reddy VT, Yang Y, Yechoor V, Mancini MA, Chan L. Berardinelli-seip congenital lipodystrophy 2/seipin is a cell-autonomous regulator of lipolysis essential for adipocyte differentiation. Mol Cell Biol 2012;32:1099–1111.

14. Cui X, Wang Y, Tang Y, Liu Y, Zhao L, Deng J, Xu G, Peng X, Ju S, Liu G, Yang H. Seipin ablation in mice results in severe generalized lipodystrophy. Hum Mol Genet 2011;20:3022–3030.

15. Prieur X, Dollet L, Takahashi M, Nemani M, Pillot B, Le May C, Mounier C, Takigawa-Imamura H, Zelenika D, Matsuda F, Feve B, Capeau J, Lathrop M, Costet P, Cariou B, Magre J. Thiazolidinediones partially reverse the metabolic disturbances observed in Bscl2/seipin-deficient mice. Diabetologia 2013;56:1813–1825.

16. Fei W, Shui G, Gaeta B, Du X, Kuerschner L, Li P, Brown AJ, Wenk MR, Parton RG, Yang H. Fld1p, a functional homologue of human seipin, regulates the size of lipid droplets in yeast. J Cell Biol 2008;180:473–482.

17. Szymanski KM, Binns D, Bartz R, Grishin NV, Li WP, Agarwal AK, Garg A, Anderson RG, Goodman JM. The lipodystrophy protein seipin is found at endoplasmic reticulum lipid droplet junctions and is important for droplet morphology. Proc Natl Acad Sci U S A 2007; 104:20890–20895.

18. Bi J, Wang W, Liu Z, Huang X, Jiang Q, Liu G, Wang Y, Huang X. Seipin promotes adipose tissue fat storage through the ER Ca(2)(+)-ATPase SERCA. Cell Metab 2014;19:861–871.

19. Ding L, Yang X, Tian H, Liang J, Zhang F, Wang G, Wang Y, Ding M, Shui G, Huang X. Seipin regulates lipid homeostasis by ensuring calcium-dependent mitochondrial metabolism. EMBO J 2018;37:e97572.

20. Yan R, Qian H, Lukmantara I, Gao M, Du X, Yan N, Yang H. Human SEIPIN Binds Anionic Phospholipids. Dev Cell 2018;47:248–256 e244.

21. Sui X, Arlt H, Brock KP, Lai ZW, DiMaio F, Marks DS, Liao M, Farese RV, Jr., Walther TC. Cryo-electron microscopy structure of the lipid droplet-formation protein seipin. J Cell Biol 2018;217:4080–4091.

22. Talukder MM, Sim MF, O’Rahilly S, Edwardson JM, Rochford JJ. Seipin oligomers can interact directly with AGPAT2 and lipin 1, physically scaffolding critical regulators of adipogenesis. Mol Metab 2015;4:199–209.

23. Sim MF, Dennis RJ, Aubry EM, Ramanathan N, Sembongi H, Saudek V, Ito D, O’Rahilly S, Siniossoglou S, Rochford JJ. The human lipodystrophy protein seipin is an ER membrane adaptor for the adipogenic PA phosphatase lipin 1. Mol Metab 2012;2:38–46.

24. Pagac M, Cooper DE, Qi Y, Lukmantara IE, Mak HY, Wu Z, Tian Y, Liu Z, Lei M, Du X, Ferguson C, Kotevski D, Sadowski P, Chen W, Boroda S, Harris TE, Liu G, Parton RG, Huang X, Coleman RA, Yang H. SEIPIN Regulates Lipid Droplet Expansion and Adipocyte Development by Modulating the Activity of Glycerol-3-phosphate Acyltransferase. Cell Rep 2016; 17:1546–1559.

25. Castro IG, Eisenberg-Bord M, Persiani E, Rochford JJ, Schuldiner M, Bohnert M. Promethin Is a Conserved Seipin Partner Protein. Cells 2019;8:268.

26. Chung J, Wu X, Lambert TJ, Lai ZW, Walther TC, Farese RV, Jr. LDAF1 and Seipin Form a Lipid Droplet Assembly Complex. Dev Cell 2019;51:551–563 e557.

27. Zhou H, Lei X, Yan Y, Lydic T, Li J, Weintraub NL, Su H, Chen W. Targeting ATGL to rescue BSCL2 lipodystrophy and its associated cardiomyopathy. JCI Insight 2019;4:e129781.

28. Chen W, Yechoor VK, Chang BH, Li MV, March KL, Chan L. The Human Lipodystrophy Gene Product BSCL2/Seipin Plays a Key Role in Adipocyte Differentiation. Endocrinology 2009; 150:4552–4561.

29. Pugach EK, Richmond PA, Azofeifa JG, Dowell RD, Leinwand LA. Prolonged Cre expression driven by the alpha-myosin heavy chain promoter can be cardiotoxic. J Mol Cell Cardiol 2015;86:54–61.

30. Zhou H, Black SM, Benson TW, Weintraub NL, Chen W. Berardinelli-Seip Congenital Lipodystrophy 2/Seipin Is Not Required for Brown Adipogenesis but Regulates Brown Adipose Tissue Development and Function. Mol Cell Biol 2016;36:2027–2038.

31. Zhou H, Lei X, Benson T, Mintz J, Xu X, Harris RB, Weintraub NL, Wang X, Chen W. Berardinelli-Seip congenital lipodystrophy 2 regulates adipocyte lipolysis, browning, and energy balance in adult animals. J Lipid Res 2015;56:1912–1925.

32. Zhou H, Xu C, Lee H, Yoon Y, Chen W. Berardinelli-Seip congenital lipodystrophy 2/SEIPIN determines brown adipose tissue maintenance and thermogenic programing. Mol Metab 2020;36:100971.

33. Ussher JR, Keung W, Fillmore N, Koves TR, Mori J, Zhang L, Lopaschuk DG, Ilkayeva OR, Wagg CS, Jaswal JS, Muoio DM, Lopaschuk GD. Treatment with the 3-ketoacyl-CoA thiolase inhibitor trimetazidine does not exacerbate whole-body insulin resistance in obese mice. J Pharmacol Exp Ther 2014;349:487–496.

34. Joubert M, Jagu B, Montaigne D, Marechal X, Tesse A, Ayer A, Dollet L, Le May C, Toumaniantz G, Manrique A, Charpentier F, Staels B, Magre J, Cariou B, Prieur X. The Sodium-Glucose Cotransporter 2 Inhibitor Dapagliflozin Prevents Cardiomyopathy in a Diabetic Lipodystrophic Mouse Model. Diabetes 2017;66:1030–1040.

35. Bai B, Yang W, Fu Y, Foon HL, Tay WT, Yang K, Luo C, Gunaratne J, Lee P, Zile MR, Xu A, Chin CWL, Lam CSP, Han W, Wang Y. Seipin Knockout Mice Develop Heart Failure With Preserved Ejection Fraction. JACC Basic Transl Sci 2019;4:924–937.

36. Liu L, Jiang Q, Wang X, Zhang Y, Lin RC, Lam SM, Shui G, Zhou L, Li P, Wang Y, Cui X, Gao M, Zhang L, Lv Y, Xu G, Liu G, Zhao D, Yang H. Adipose-specific knockout of SEIPIN/BSCL2 results in progressive lipodystrophy. Diabetes 2014;63:2320–2331.

37. Han S, Binns DD, Chang YF, Goodman JM. Dissecting seipin function: the localized accumulation of phosphatidic acid at ER/LD junctions in the absence of seipin is suppressed by Sei1p(DeltaNterm) only in combination with Ldb16p. BMC Cell Biol 2015;16:29.

38. Wolinski H, Hofbauer HF, Hellauer K, Cristobal-Sarramian A, Kolb D, Radulovic M, Knittelfelder OL, Rechberger GN, Kohlwein SD. Seipin is involved in the regulation of phosphatidic acid metabolism at a subdomain of the nuclear envelope in yeast. Biochim Biophys Acta 2015; 1851:1450–1464.

39. Wang H, Becuwe M, Housden BE, Chitraju C, Porras AJ, Graham MM, Liu XN, Thiam AR, Savage DB, Agarwal AK, Garg A, Olarte MJ, Lin Q, Frohlich F, Hannibal-Bach HK, Upadhyayula S, Perrimon N, Kirchhausen T, Ejsing CS, Walther TC, Farese RV. Seipin is required for converting nascent to mature lipid droplets. Elife 2016;5:e16582.

40. Banke NH, Wende AR, Leone TC, O’Donnell JM, Abel ED, Kelly DP, Lewandowski ED. Preferential oxidation of triacylglyceride-derived fatty acids in heart is augmented by the nuclear receptor PPARalpha. Circ Res 2010;107:233–241.

41. Lehman JJ, Boudina S, Banke NH, Sambandam N, Han X, Young DM, Leone TC, Gross RW, Lewandowski ED, Abel ED, Kelly DP. The transcriptional coactivator PGC-1alpha is essential for maximal and efficient cardiac mitochondrial fatty acid oxidation and lipid homeostasis. Am J Physiol Heart Circ Physiol 2008;295:H185–196.

42. Kolwicz SC, Jr., Olson DP, Marney LC, Garcia-Menendez L, Synovec RE, Tian R. Cardiac-specific deletion of acetyl CoA carboxylase 2 prevents metabolic remodeling during pressure-overload hypertrophy. Circ Res 2012;111:728–738.

43. Ghosh M, Niyogi S, Bhattacharyya M, Adak M, Nayak DK, Chakrabarti S, Chakrabarti P. Ubiquitin Ligase COP1 Controls Hepatic Fat Metabolism by Targeting ATGL for Degradation. Diabetes 2016;65:3561–3572.

44. Boudina S, Sena S, Theobald H, Sheng X, Wright JJ, Hu XX, Aziz S, Johnson JI, Bugger H, Zaha VG, Abel ED. Mitochondrial energetics in the heart in obesity-related diabetes: direct evidence for increased uncoupled respiration and activation of uncoupling proteins. Diabetes 2007;56:2457–2466.

45. Kiebish MA, Yang K, Liu X, Mancuso DJ, Guan S, Zhao Z, Sims HF, Cerqua R, Cade WT, Han X, Gross RW. Dysfunctional cardiac mitochondrial bioenergetic, lipidomic, and signaling in a murine model of Barth syndrome. J Lipid Res 2013;54:1312–1325.

46. Finck BN, Lehman JJ, Leone TC, Welch MJ, Bennett MJ, Kovacs A, Han X, Gross RW, Kozak R, Lopaschuk GD, Kelly DP. The cardiac phenotype induced by PPARalpha overexpression mimics that caused by diabetes mellitus. J Clin Invest 2002;109:121–130.

47. Mihalik SJ, Goodpaster BH, Kelley DE, Chace DH, Vockley J, Toledo FG, DeLany JP. Increased levels of plasma acylcarnitines in obesity and type 2 diabetes and identification of a marker of glucolipotoxicity. Obesity (Silver Spring) 2010;18:1695–1700.

48. Ebeling P, Tuominen JA, Arenas J, Garcia Benayas C, Koivisto VA. The association of acetyl-L-carnitine with glucose and lipid metabolism in human muscle in vivo: the effect of hyperinsulinemia. Metabolism 1997;46:1454–1457.

49. Dyck JR, Cheng JF, Stanley WC, Barr R, Chandler MP, Brown S, Wallace D, Arrhenius T, Harmon C, Yang G, Nadzan AM, Lopaschuk GD. Malonyl coenzyme a decarboxylase inhibition protects the ischemic heart by inhibiting fatty acid oxidation and stimulating glucose oxidation. Circ Res 2004;94:e78–84.

50. Kantor PF, Lucien A, Kozak R, Lopaschuk GD. The antianginal drug trimetazidine shifts cardiac energy metabolism from fatty acid oxidation to glucose oxidation by inhibiting mitochondrial long-chain 3-ketoacyl coenzyme A thiolase. Circ Res 2000;86:580–588.

51. Liu Z, Chen JM, Huang H, Kuznicki M, Zheng S, Sun W, Quan N, Wang L, Yang H, Guo HM, Li J, Zhuang J, Zhu P. The protective effect of trimetazidine on myocardial ischemia/reperfusion injury through activating AMPK and ERK signaling pathway. Metabolism 2016;65:122–130.

52. Fragasso G, Palloshi A, Puccetti P, Silipigni C, Rossodivita A, Pala M, Calori G, Alfieri O, Margonato A. A randomized clinical trial of trimetazidine, a partial free fatty acid oxidation inhibitor, in patients with heart failure. J Am Coll Cardiol 2006;48:992–998.

53. Lee L, Campbell R, Scheuermann-Freestone M, Taylor R, Gunaruwan P, Williams L, Ashrafian H, Horowitz J, Fraser AG, Clarke K, Frenneaux M. Metabolic modulation with perhexiline in chronic heart failure: a randomized, controlled trial of short-term use of a novel treatment. Circulation 2005; 112:3280–3288.

54. Tuunanen H, Engblom E, Naum A, Nagren K, Scheinin M, Hesse B, Juhani Airaksinen KE, Nuutila P, Iozzo P, Ukkonen H, Opie LH, Knuuti J. Trimetazidine, a metabolic modulator, has cardiac and extracardiac benefits in idiopathic dilated cardiomyopathy. Circulation 2008; 118:1250–1258.

55. Luptak I, Balschi JA, Xing Y, Leone TC, Kelly DP, Tian R. Decreased contractile and metabolic reserve in peroxisome proliferator-activated receptor-alpha-null hearts can be rescued by increasing glucose transport and utilization. Circulation 2005; 112:2339–2346.

56. Li H, Ma Z, Zhai Y, Lv C, Yuan P, Zhu F, Wei L, Li Q, Qi X. Trimetazidine Ameliorates Myocardial Metabolic Remodeling in Isoproterenol-Induced Rats Through Regulating Ketone Body Metabolism via Activating AMPK and PPAR alpha. Front Pharmacol 2020;11:1255.

57. Di Napoli P, Di Giovanni P, Gaeta MA, D’Apolito G, Barsotti A. Beneficial effects of trimetazidine treatment on exercise tolerance and B-type natriuretic peptide and troponin T plasma levels in patients with stable ischemic cardiomyopathy. Am Heart J 2007; 154:602 e601–605.

58. Liu F, Yin L, Zhang L, Liu W, Liu J, Wang Y, Yu B. Trimetazidine improves right ventricular function by increasing miR-21 expression. Int J Mol Med 2012;30:849–855.

59. Stanley WC, Dabkowski ER, Ribeiro RF, Jr., O’Connell KA. Dietary fat and heart failure: moving from lipotoxicity to lipoprotection. Circ Res 2012;110:764–776.

60. Agah R, Frenkel PA, French BA, Michael LH, Overbeek PA, Schneider MD. Gene recombination in postmitotic cells. Targeted expression of Cre recombinase provokes cardiac-restricted, site-specific rearrangement in adult ventricular muscle in vivo. J Clin Invest 1997; 100:169–179.

61. Xu W, Zhou H, Xuan H, Saha P, Wang G, Chen W. Novel metabolic disorders in skeletal muscle of Lipodystrophic Bscl2/Seipin deficient mice. Mol Cell Endocrinol 2019;482:1–10.

62. Ackers-Johnson M, Li PY, Holmes AP, O’Brien SM, Pavlovic D, Foo RS. A Simplified, Langendorff-Free Method for Concomitant Isolation of Viable Cardiac Myocytes and Nonmyocytes From the Adult Mouse Heart. Circ Res 2016; 119:909–920.

63. Bray MS, Shaw CA, Moore MW, Garcia RA, Zanquetta MM, Durgan DJ, Jeong WJ, Tsai JY, Bugger H, Zhang D, Rohrwasser A, Rennison JH, Dyck JR, Litwin SE, Hardin PE, Chow CW, Chandler MP, Abel ED, Young ME. Disruption of the circadian clock within the cardiomyocyte influences myocardial contractile function, metabolism, and gene expression. Am J Physiol Heart Circ Physiol 2008;294:H1036–1047.

64. Tsai JY, Kienesberger PC, Pulinilkunnil T, Sailors MH, Durgan DJ, Villegas-Montoya C, Jahoor A, Gonzalez R, Garvey ME, Boland B, Blasier Z, McElfresh TA, Nannegari V, Chow CW, Heird WC, Chandler MP, Dyck JR, Bray MS, Young ME. Direct regulation of myocardial triglyceride metabolism by the cardiomyocyte circadian clock. J Biol Chem 2010;285:2918–2929.

