## Supplemental data for "BSCL2/Seipin Deficiency in Heart Causes Energy Deficit and Heart Failure via Inducing Excessive Lipid Catabolism"

**Supplemental Materials and Methods**

**Histology and trichrome staining**

Tissues were formalin fixed, paraffin embedded, sectioned and stained with Hematoxylin–eosin (H&E). Trichrome staining was performed using Trichrome Staining Kit (Abcam, ab150686) according to manufacturers’ directions.

**Plasma biochemistry**

Blood glucose levels were measured by One-touch Ultra glucose meter. Glycerol and NEFA levels were determined using a free glycerol reagent (Sigma-Aldrich) and WAKO NEFA analysis kit (NEFA-HR(2); Wako Pure Chemical Industries), respectively. Plasma triglyceride and cholesterol levels were measured by colorimetrical analyses using triglyceride assay kit (Infinity^TM^ triglycerides kit, Thermo Fisher Scientific) and total cholesterol assay kit (Infinity^TM^ triglycerides kit, Thermo Fisher Scientific) respectively.

**Measurement of reactive oxygen species**

Fluorescence of reaction oxygen species (ROS) were detected by incubating frozen sections with 5 μM DCFDA for 30 min. Fluorescence intensity was first normalized to protein level and ROS activities were expressed as fold changes relative to Ctrl mice. MDA, an indicator of lipid peroxidation as a part of thiobarbituric acid reacting substances (TBARS), was assayed using TBARS Assay Kit (Caymen Chemicals) as instructed.

**Mitochondrial isolation and measurement of mitochondrial respiration**

For Seahorse bioenergetics analysis, fresh ventricles were minced in ice-cold fiber relaxation buffer (KCl 100 mM, EGTA 5 mM, HEPES 5 mM, pH 7.0) and homogenized in a glass douncer followed by differential centrifugation using ice-cold HES buffer (HEPES 5 mM, EDTA 1 mM, Sucrose 0.25 M, pH 7.4). Mitochondrial protein yield was determined by Bradford assay and 3 μg mitochondria were seeded per well by centrifugation. Mitochondrial respiration was assessed in respiration buffer (220 mM mannitol, 70 mM sucrose, 10 mM KH2PO4, 5 mM MgCl2, 2 mM HEPES, 1.0 mM EGTA, 1 mM EDTA, 0.2 % BSA, pH 7.2) containing 10 mM succinate and 2 μM rotenone with sequentially addition of 4 mM adenosine 5’-diphosphate (ADP) (Complex V substrate), 2.5 μg/ml oligomycin (Complex V inhibitor), 4 μM carbonyl cyanide 4-(trifluoromethoxy)phenylhydrazone (FCCP) (mitochondrial uncoupler for maximal respiration), and 4 μM antimycin A (Complex III inhibitor). Oxygen consumption rates (OCRs) were measured using an XF24 Extracellular Flux Analyzer (Seahorse Bioscience) as previously described^1^. Coupled respiration was the oligomycin-sensitive OCR while uncoupled was the OCR difference between oligomycin and antimycin A additions.

**Mouse heart mitochondria DNA content**

Total DNA was extracted from ventricles and mitochondrial DNA (mtDNA) content was analyzed by RT-PCR of mtDNA-encoded 16s RNA (*MT-Rnr2*) to nuclear DNA (nDNA)-encoded hexokinase 2 (*Hk2*) intron 9 as described previously^2^.

**Tissue glycogen measurement**

Ventricle glycogen content was measured by homogenizing tissues in 0.5 N KOH. Glycogen was then precipitated using ethanol and digested with amyloglycosidase (Sigma). The released glucose concentrations were then quantified using a glucose hexokinase assay kit (Thermo Fisher Scientific) and normalized to tissue weights as previously described^3^.

**Antibody information**

The following antibodies were used: rabbit antibodies against Phospho-PKA substrate (9624), HSL (4107), ATGL (2138), Phosphor-Phospholamban (Ser16/Thr17) (8496), Phospholamban (8495), HSP60 (12165) are from Cell Signaling Technology. CD36 (18836-1-AP), GAPDH (60004-1-IG), CPT1β (22170-1-AP), Prohibitin (10787-1-AP), SOD1 (10269-1-AP), SOD2 (24127-1-AP) and Catalase (21260-1-AP) are from Proteintech. Total OXPHOS Rodent WB Antibody Cocktail (ab110413) and PPARα (ab24509) are from AbCam.

**Supplemental Tables**

**Supplemental Table S1. Echocardiography in mice under normal chow diet.**

| **Age** | **3 month-old** | | | **6 month-old** | | |
| --- | --- | --- | --- | --- | --- | --- |
| **Genotype** | **Ctrl (n=8)** | ***Cre+; Bscl2^w/w^* (n=8)** | **cKO (n=12)** | **Ctrl (n=11)** | ***Cre+; Bscl2^w/w^* (n=8)** | **cKO (n=9)** |
| **Body weights (g)** | 24.5 ± 0.6 | 26.9 ± 0.4 | 25.2 ± 0.4 | 29.5 ± 0.6 | 32.1 ± 1.74 | 30.6 ± 0.6 |
| **LVPWd (mm)** | 0.74 ± 0.02 | 0.72 ± 0.06 | 0.75 ± 0.04 | 0.86 ± 0.05 | 0.92 ± 0.03 | 0.79 ± 0.04* |
| **LVAWd (mm)** | 0.71 ± 0.03 | 0.8 ± 0.03 | 0.76 ± 0.03 | 0.82 ± 0.02 | 0.75 ± 0.04 | 0.69 ± 0.06* |
| **LVIDd (mm)** | 3.55 ± 0.07 | 3.61 ± 0.07 | 3.59 ± 0.01 | 3.68 ± 0.06 | 3.9 ± 0.12 | 4.1 ± 0.12** |
| 3 and 6-month old male *Cre-; Bscl2^f/f^* (Ctrl), *Cre+; Bscl2^w/w^* and *Cre+; Bscl2^f/f^* (*Bscl2^cKO^*, simplified as cKO) mice were kept under normal chow diet. LVPWd: left ventricle post wall thickness at end diastole; LVAWd: left ventricle post wall anterior wall thickness at end diastole; LVIDd: left ventricle internal diameter at end diastole. *: *P*< 0.05; **: *P*< 0.005 vs Ctrl mice within the same age group. One-way ANOVA with Dunnett’s multiple comparisons test. | | | | | | |

**Supplemental Table S2. Plasma parameters and echocardiography in mice fed with normal chow and high fat diets.**

| **Diet** | **NCD** | | **HFD** | |
| --- | --- | --- | --- | --- |
| **Genotype** | **Ctrl** | **cKO** | **Ctrl** | **cKO** |
| **Body weight (g)** | 32.1 ± 0.6 | 31.4 ± 0.5 | 43.7 ± 1.2^##^ | 45.7 ± 0.7^##^ |
| **Glucose (mg/dL)** | 153 ± 5 | 152 ± 4 | 233 ± 8^##^ | 221 ± 10^##^ |
| **TG (mg/dL)** | 59.2 ± 4.9 | 74.1 ± 3.9 | 53.2 ± 1.2 | 61.9 ± 1.7 |
| **TC (mg/dL)** | 82± 9.2 | 64.4 ± 6.3 | 207.5 ± 10.8^##^ | 203.7 ± 7.8^##^ |
| **NEFA (mM)** | 0.49 ± 0.07 | 0.35 ± 0.02 | 1.38 ± 0.12^##^ | 1.42 ± 0.1^##^ |
| **Glycerol (mg/dL)** | 19.8 ± 1.9 | 25.3 ± 2.9 | 47.1 ± 4^#^ | 43.8 ± 3.4^#^ |
| **LVPWd (mm)** | 0.83 ± 0.06 | 0.76 ± 0.04 | 0.92 ± 0.06 | 0.80 ± 0.03^#^ |
| **LVAWd (mm)** | 0.77 ± 0.04 | 0.68 ± 0.05 | 0.95 ± 0.03^#^ | 0.83 ± 0.02^#^ |
| **LVAWs (mm)** | 1.4 ± 0.04 | 1.15 ± 0.05** | 1.55 ± 0.05 | 1.39 ± 0.03*^##^ |
| **LVIDd (mm)** | 3.84 ± 0.07 | 4.11 ± 0.12 | 3.84 ± 4 | 4.14 ± 0.08 |
| **EF (%)** | 72.6 ± 0.75 | 56.8 ± 1.13** | 70.4 ± 1.42 | 67.9 ± 1.0**^##^ |
| Body weights, plasma biochemistry and echocardiography were analyzed in ad-libitum 6 month-old male *Cre-; Bscl2^f/f^* (Ctrl) and *Bscl2^cKO^* (cKO) mice fed with normal chow diet (NCD) or high fat diet (HFD) starting at 3 months old for additional 3 months. TG: triglyceride; TC: total cholesterol; NEFA: nonesterified fatty acid. TG: triglyceride; TC: total cholesterol; NEFA: nonesterified fatty acid. LVPWd: left ventricle post wall thickness at end diastole; LVAWd: left ventricle post wall anterior wall thickness at end diastole; LVAWs: left ventricle post wall anterior wall thickness at end systole; LVIDd: left ventricle internal diameter at end diastole; EF: ejection fraction. Data were presented as means ± SEM. For plasma parameters, NCD-Ctrl, *n*=9; NCD-cKO, *n*=9; HFD-Ctrl, *n*=7; HFD-cKO, *n*=11. For echocardiography, NCD-Ctrl, *n*=9; NCD-cKO, *n*=12; HFD-Ctrl, *n*=12; HFD-cKO, *n*=16. *P*<0.005 vs PBS treated same genotype. Two-way ANOVA with Turkey’s multiple comparisons tests. | | | | |

**Supplemental Table 3. Plasma parameters and echocardiography in mice treated with trimetazidine.**

| **Treatment** | **PBS** | | **TMZ** | |
| --- | --- | --- | --- | --- |
| **Genotype** | **Ctrl (n=6)** | **cKO (n=9)** | **Ctrl (n=6)** | **cKO (n=9)** |
| **Glucose (mg/dL)** | 116 ± 3.4 | 116.3 ± 5.2 | 122.6 ± 4.3 | 128.2 ± 5.2 |
| **TG (mg/dL)** | 81 ± 7.5 | 78.5 ± 5.6 | 75.7 ± 5 | 69.5 ± 3.6 |
| **TC (mg/dL)** | 71 ± 7.1 | 61.3 ± 7.1 | 90.2 ± 6.2^#^ | 78.6 ± 3.5^#^ |
| **NEFA (mM)** | 0.76 ± 0.24 | 0.66 ± 0.12 | 0.6 ± 0.13 | 0.49 ± 0.08 |
| **Glycerol (mg/dL)** | 13.7 ± 0.9 | 14.3 ± 0.9 | 14.4 ± 0.8 | 15 ± 0.9 |
| **LVPWd (mm)** | 0.67 ± 0.06 | 0.75 ± 0.05 | 0.81 ± 0.05 | 0.63 ± 0.03 |
| **LVAWd (mm)** | 0.79 ± 0.05 | 0.71 ± 0.05 | 0.80 ± 0.03 | 0.78 ± 0.04^#^ |
| **LVAWs (mm)** | 1.49 ± 0.06 | 1.11 ± 0.06** | 1.44 ± 0.03 | 1.27 ± 0.04 |
| **LVIDd (mm)** | 3.75 ± 0.06 | 4.12 ± 0.10* | 3.82 ± 0.15 | 4.11 ± 0.05 |
| **EF (%)** | 73.78 ± 1.62 | 52.5 ± 1.92** | 72.38 ± 1.57 | 60.9 ± 0.83**^##^ |
| Plasma biochemistry and echocardiography were analyzed in ad-libitum 7.5 month-old male *Cre-; Bscl2^f/f^* (Ctrl) and *Bscl2^cKO^* (cKO) mice i.p. injected with PBS or trimetazidine (TMZ) for 6 weeks starting at 6 months old. TG: triglyceride; TC: total cholesterol; NEFA: nonesterified fatty acid. LVPWd: left ventricle post wall thickness at end diastole; LVAWd: left ventricle post wall anterior wall thickness at end diastole; LVAWs: left ventricle post wall anterior wall thickness at end systole; LVIDd: left ventricle internal diameter at end diastole; EF: ejection fraction. Data were presented as means ± SEM. *: *P*<0.05; **: *P*<0.005 vs Ctrl mice under the same treatment. #: *P*<0.05; ##: *P*<0.005 vs PBS treated same genotype. Two-way ANOVA with Turkey’s multiple comparisons tests. | | | | |

**Supplemental Table S4. Quantitative real-time PCR primer sequences for murine genes**

| **Gene name** |  | **Primer sequence** | **Gene name** |  | **Primer sequence** |
| --- | --- | --- | --- | --- | --- |
| ***36B4*** | 5F | CGCTTTCTGGAGGGTGTCCGC | ***Lpl*** | 5F | CCTAAGGACCCCTGAAGAC |
|  | 3R | TGCCAGGACGCGCTTGTACC |  | 3R | GACATTGGAGTCAGGTTCTC |
| ***Acadl*** | 5F | AGCCTCCGTGGAGTTGCACA | ***Myh6*** | 5F | CGCCTATGAGGAGTCTCTGG |
|  | 3R | CCAGGAACTACGTGAAGCAAAG |  | 3R | TTCTCCACCTCCAGCTGTTT |
| ***Actb*** | 5F | GACGGCCAGGTCATCACTAT | ***Myh7*** | 5F | CGCCTATGAGGAGTCTCTGG |
|  | 3R | CTTCTGCATCCTGTCAGCAA |  | 3R | TCCAGTTGCTTTCGGATCTT |
| ***Bscl2*** | 5F | GCTCTTCTGCACCATCCTTC | ***Nppa*** | 5F | ATTGACAGGATTGGAGCCCAGAGT |
|  | 3R | CGGTGGAGGAATCACAGTC |  | 3R | TGACACACCACAAGGGCTTAGGAT |
| ***Cd36*** | 5F | CGTTTCAACTCTCACACACATAAG | ***Nppb*** | 5F | ATCTCCTGAAGGTGCTGTCC |
|  | 3R | TGAGACTCTGAAAGGATCAGCA |  | 3R | AGCTGTCTCTGGGCCATTT |
| ***Acox1*** | 5F | CCTGACAGAAGCCTACAAG | ***Pdk4*** | 5F | CTCCTTCGGTGCAGCTGG |
|  | 3R | TGTCTTGAATCTTGGGGAGTT |  | 3R | GTCCACTGTGCAGGTGTCT |
| ***Cpt1β*** | 5F | TTGCCCTACAGCTGGCTCATTTCC | ***Pgc1α*** | 5F | CCCTGCCATTGTTAAGACC |
|  | 3R | GCACCCAGATGATTGGGATACTGT |  | 3R | TGCTGCTGTTCCTGTTTTC |
| ***Col1a2*** | 5F | GTCCTAGTCGATGGCTGCTC | ***Pnpla2*** | 5F | GATGTGCAAACAGGGCTACA |
|  | 3R | GTCAGCACCACCAATGTCC |  | 3R | CTTCCTCTGCATCCTCTTCC |
| ***Gdf15*** | 5F | AGCCGAGAGGACTCGAAC | ***Pparα*** | 5F | CCACGAAGCCTACCTGAAGA |
|  | 3R | GTTGACGCGGAGTAGCAG |  | 3R | ACTGGCAGCAGTGGAAGAAT |
| ***Glut1*** | 5F | GCTTTGTGGCCTTCTTTGAA | ***Ppia*** | 5F | CTGTTTGCAGACAAAGTTCCA |
|  | 3R | AAGAAGAGCACGAGGAGCAC |  | 3R | AGGATGAAGTTCTCATCCTCA |
| ***Glut4*** | 5F | CGGCTCTGACGATGGGGA | ***Tfam*** | 5F | CCACAGAACAGCTACCCAAA |
|  | 3R | GGTGCCTTGTGGGATGGA |  | 3R | CATCAGCTGACTTGGAGTTA |
| ***Lipe*** | 5F | GCTCTTCTTCGAGGGTGATG |  |  |  |
|  | 3R | ACACTGAGGCCTGTCTCGTT |  |  |  |

**Supplemental Figures**

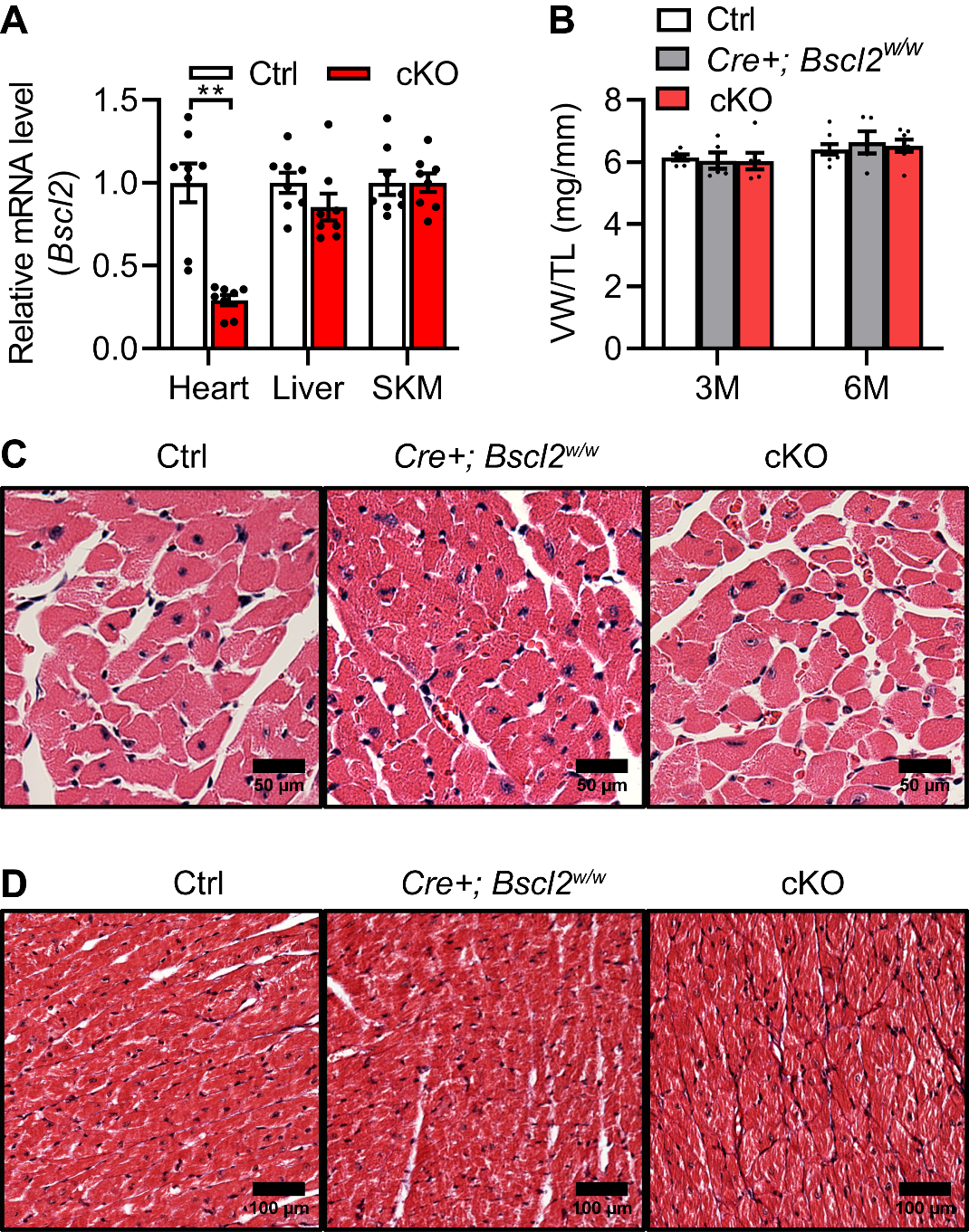

**Supplemental Figure S1. Mice with cardiac-specific deletion of BSCL2 develop dilated cardiomyopathy.**

**(A)** RT-PCR analysis of *Bscl2* gene expression in ventricles from 3 month-old male *Cre-*; *Bscl2^f/f^* (Ctrl), *Cre+*; *Bscl2^w/w^*, and *Cre+*; *Bscl2^f/f^* (cKO) mice. *n*=8 per group. **: *P*< 0.005 vs Ctrl, unpaired t tests. **(B)** Ventricle weight (VW) normalized to tibia length (TL) in 3 month-old (3M) and 6 month-old (6M) male Ctrl, *Cre+*; *Bscl2^w/w^* and cKO mice. 3M old: Ctrl, *n*=8; *Cre+*; *Bscl2^w/w^*, *n*=8; cKO, *n*=12. 6M old: Ctrl, *n*=11; *Cre+*; *Bscl2^w/w^*, *n*=8; cKO, *n*=9. **(C)** Representative images of hematoxylin-eosin staining of left ventricle apexes in male 6 month-old mice. Scale bar = 50 µm. **(D)** Histochemical assessment of fibrosis (trichrome staining) in male 6 months-old mice. Scale bar = 100 µm.

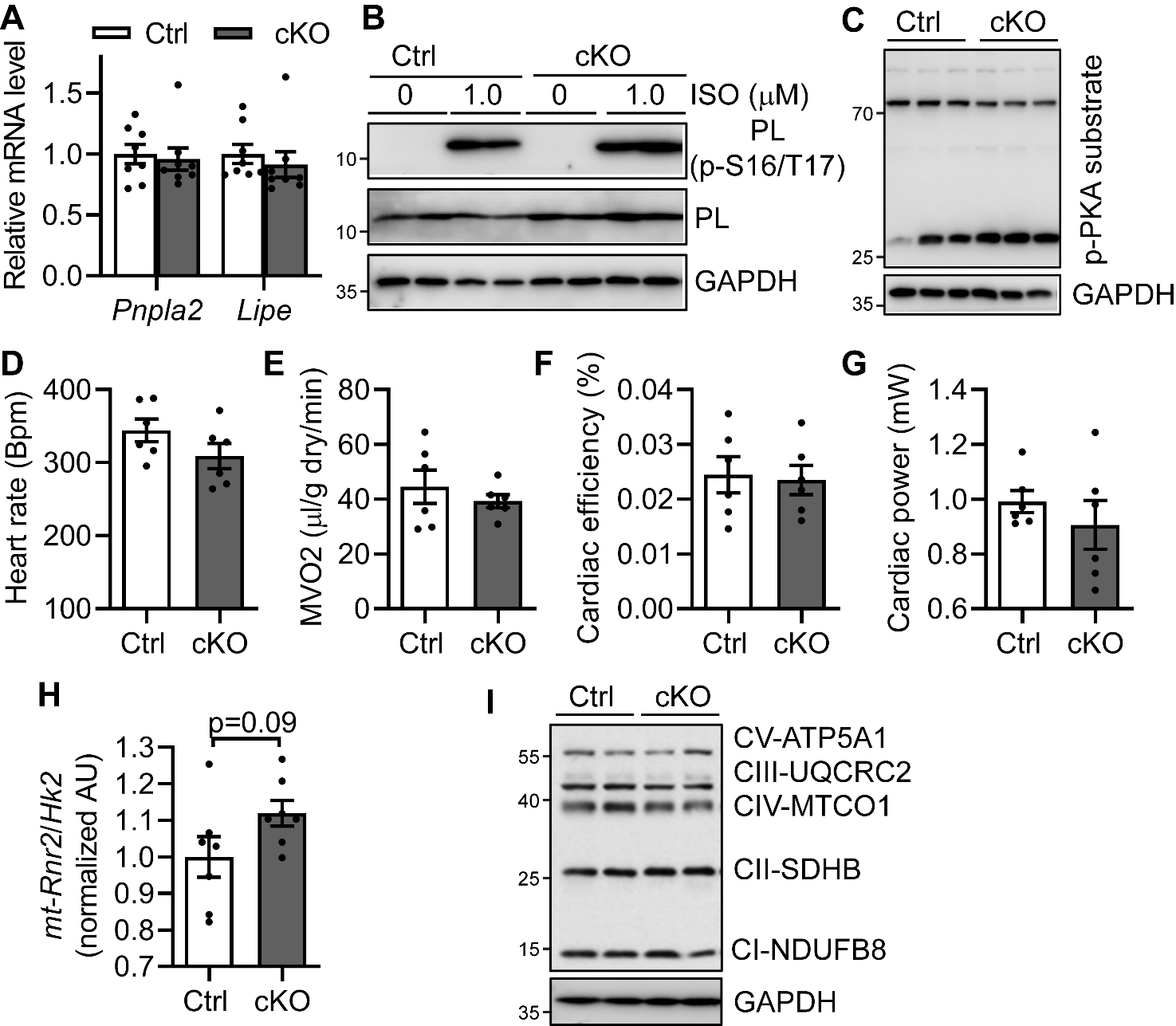

**Supplemental Figure S2. Cardiac–specific deletion of BSCL2 regulates cardiac ATGL expression and substrate metabolism.**

**(A)** mRNA expression of lipolytic genes in 3-month-old *Cre-*; *Bscl2^f/f^* (Ctrl) and *Bscl2^cKO^* (cKO) mice. *n*=8 per group. **(B)** Representative Western blotting to demonstrate phosphorylation of phospholamban (PL) 20 min after addition of 1 µM isoproterenol (ISO) in adult cardiomyocytes isolated from 3-month-old Ctrl and cKO mice. Two independent experiments. **(C)** Basal PKA-mediated substrate phosphorylation in homogenates from 3-month-old Ctrl and cKO ventricles. *n*=3 per group. Three independent experiments. **(D)** Heart rate; **(E)** myocardial oxygen consumption (MVO2); **(F)** cardiac efficiency; and **(G)** cardiac power in *ex vivo* perfused working hearts. **(D-G)**: *n*=6 per group. **(H)** Relative mtDNA content in ventricles assessed by RT-PCR and calculated from copy numbers of the mtDNA-encoded *mt-Rnr2* gene and the nuclear DNA-encoded *Hk2* intron 9 gene. Data were presented as fold change compared with Ctrl, arbitrarily defined as 1. *n*=7 per group. **(I)** Representative Western blotting of mitochondrial complex proteins in ventricles of 3-month-old Ctrl and cKO mice. *n*=2 per group. Three independent experiments. *: *P*< 0.05 with unpaired t test (parametric).

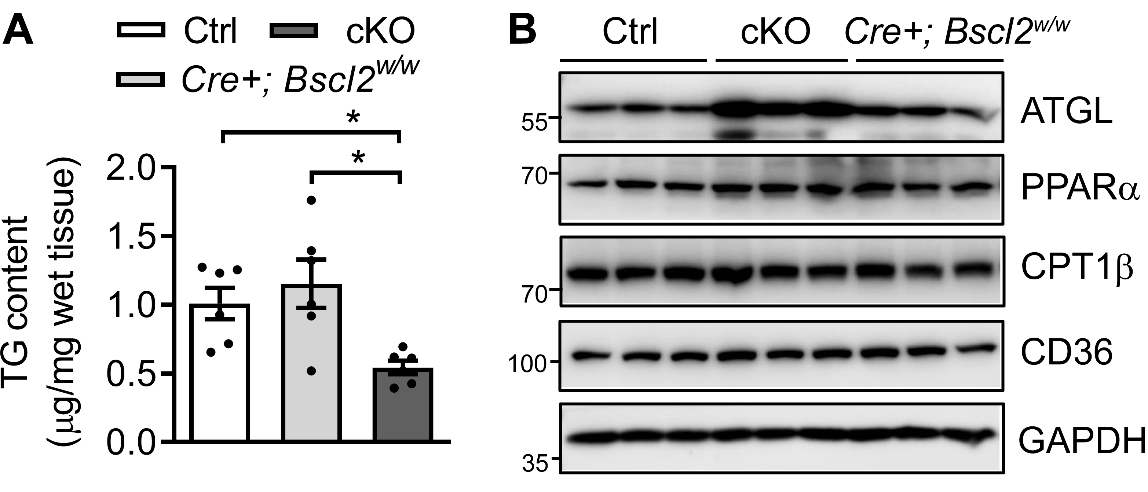

**Supplemental Figure S3. 6 month-old *Bscl2^cKO^* mice Exhibited cardiac lipid remodeling.**

**(A)** Enzymatic quantification of ventricular TG contents, *n*=6 per group, and **(B)** representative Western blotting in ventricles of 6-month-old *Cre-*; *Bscl2^f/f^* (Ctrl), *Bscl2^cKO^* (cKO) mice and *Cre+; Bscl2^w/w^* mice. *n*=3 per group. Two independent experiments. *: *P*< 0.05, One-way ANOVA followed by Dunnett’s multiple comparisons test.

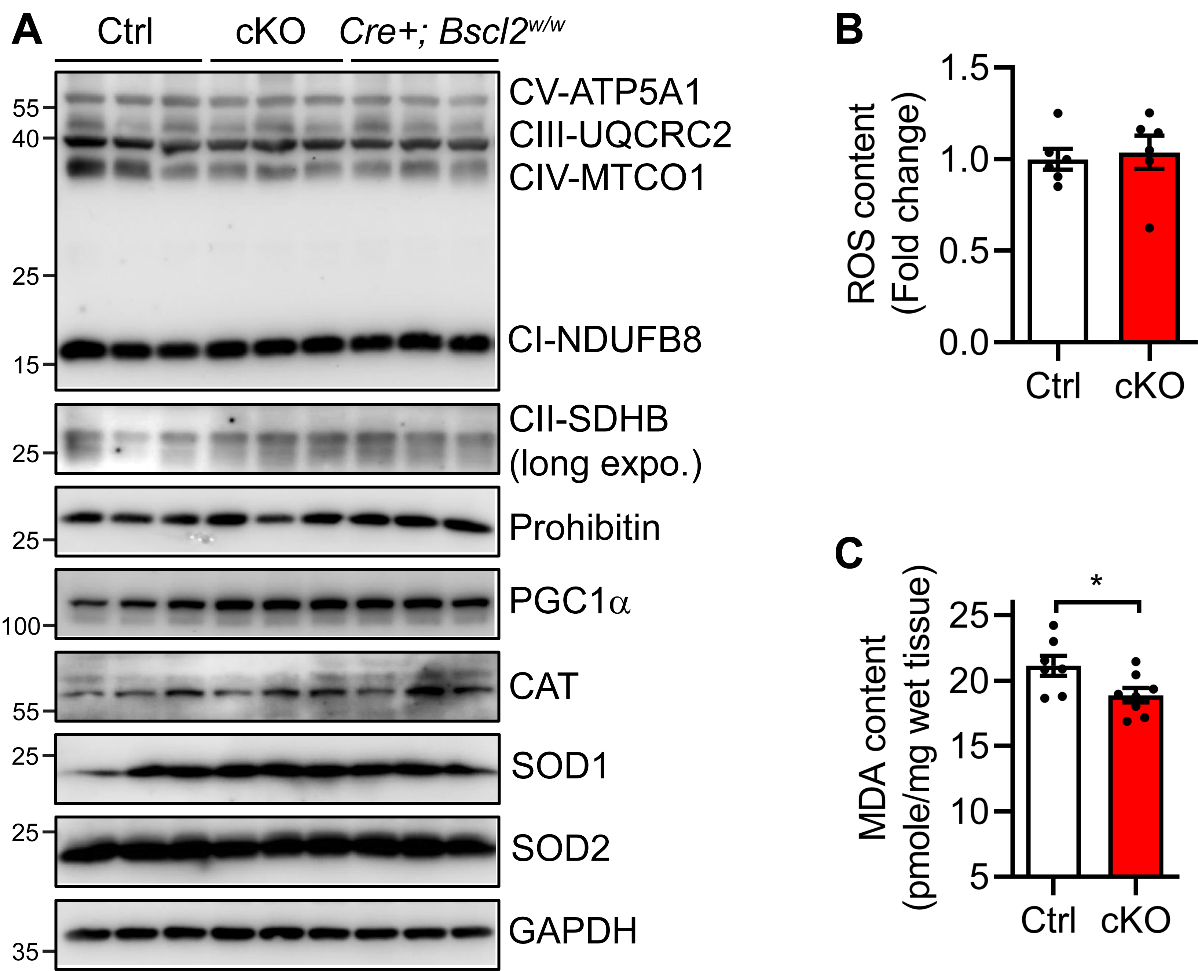

**Supplemental Figure S4. Cardiac dysfunction in *Bscl2^cKO^* mice is not associated with mitochondrial dysfunction and oxidative stress.**

**(A)** Representative Western blotting in ventricles of 6-month-old *Cre-*; *Bscl2^f/f^* (Ctrl), *Bscl2^cKO^* (cKO) mice and *Cre+; Bscl2^w/w^* mice. *n*=3 per group. Two independent experiments. **(B)** Ventricular ROS contents as measured by incubating ventricular homogenates with DCFDA dye. Data were presented as fold change with Ctrl normalized to 1. *n*=6 per group. **(C)** Ventricular MDA contents as measured by TBARS kit. Ctrl, *n*=7; cKO, *n*=8. *: *P*< 0.05, unpaired t tests (parametric).

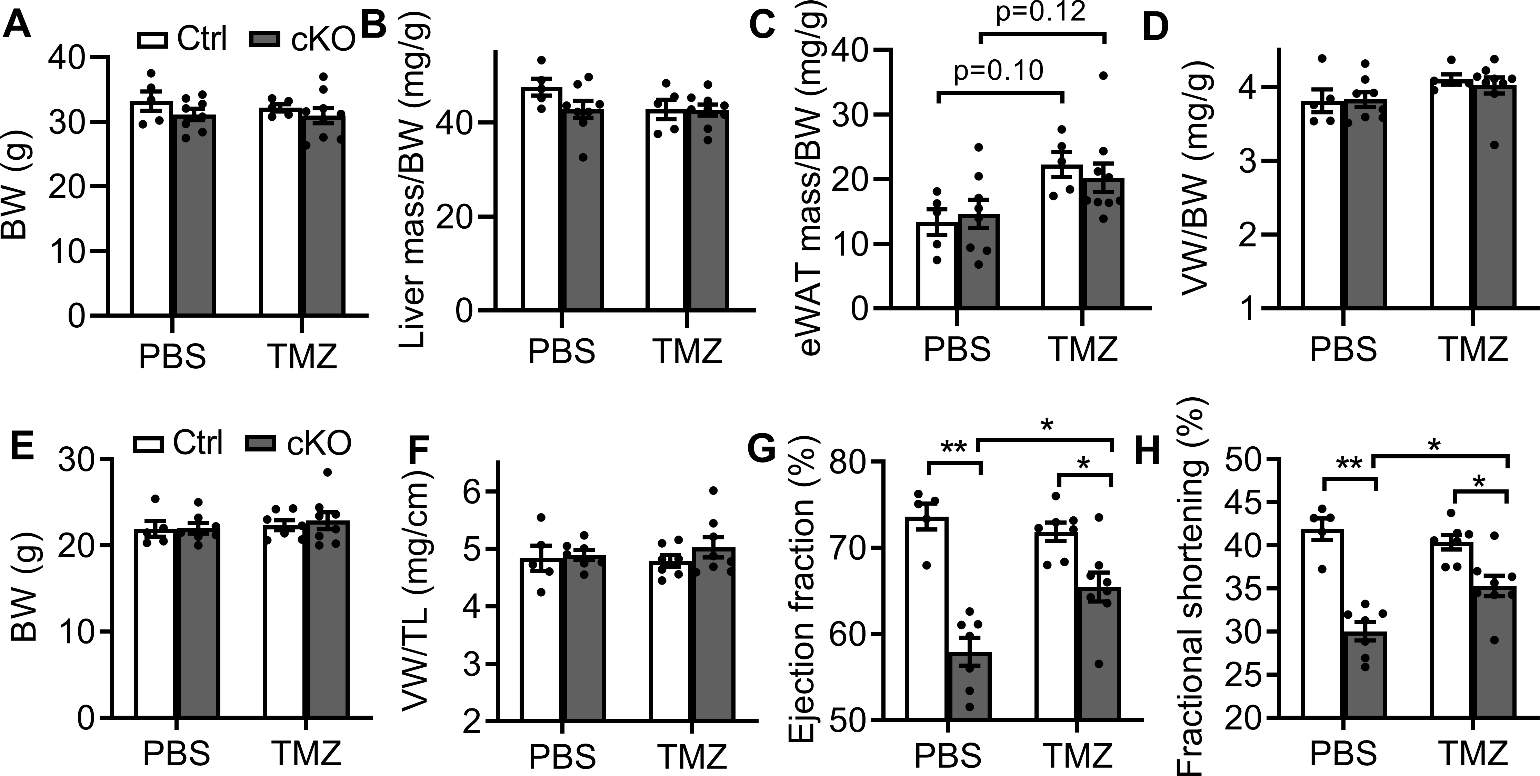

**Supplemental Figure S5. Inhibition of fatty acid oxidation partially rescues cardiac function in *Bscl2^cKO^* mice.**

6-month-old male *Cre-*; *Bscl2^f/f^* (Ctrl) and *Bscl2^cKO^* (cKO) mice were daily i.p. injected with PBS or trimetazidine (TMZ) at 15 mg/kg body weights (BW) for a total of 6 weeks. **(A)** Body weight (BW); **(B)** ratio of liver mass to BW; **(C)** ratio of epididymal white fat (eWAT) to BW and **(D)** ratio of ventricle weight (VW) to BW at the end of treatment. For **A-D**, PBS-Ctrl, *n*=5; PBS-cKO, *n*=8. TMZ-Ctrl, *n*=5, TMZ-cKO, *n*=9. **(E-H)** 6 month-old female Ctrl and cKO mice were treated with TMZ as above. **(E)** BW; **(F)** ratio of ventricle weight (VW) to tibia length (TL); **(G)** ejection fraction and **(H)** fractional shortening were assessed after 6 weeks of PBS or TMZ injection. For **E-H**, PBS-Ctrl, *n*=5; PBS-cKO, *n*=7. TMZ-Ctrl, *n*=7, TMZ-cKO, *n*=8. *: *P*< 0.05; **: *P*< 0.005. Two-way ANOVA followed by Tukey’s post-hoc tests.

**Supplemental References**

1.	Zhou H, Xu C, Lee H, Yoon Y, Chen W. Berardinelli-Seip congenital lipodystrophy 2/SEIPIN determines brown adipose tissue maintenance and thermogenic programing. *Mol Metab* 2020;**36**:100971.

2.	Zhou H, Black SM, Benson TW, Weintraub NL, Chen W. Berardinelli-Seip Congenital Lipodystrophy 2/Seipin Is Not Required for Brown Adipogenesis but Regulates Brown Adipose Tissue Development and Function. *Mol Cell Biol* 2016;**36**:2027-2038.

3.	Chen W, Zhou H, Saha P, Li L, Chan L. Molecular mechanisms underlying fasting modulated liver insulin sensitivity and metabolism in male lipodystrophic Bscl2/Seipin-deficient mice. *Endocrinology* 2014;**155**:4215-4225.
